# Distinct pathways of homologous recombination controlled by the SWS1-SWSAP1-SPIDR complex

**DOI:** 10.1101/2020.05.15.098848

**Authors:** Rohit Prakash, Thomas Sandoval, Florian Morati, Jennifer A Zagelbaum, Brett Taylor, Raymond Wang, Emilie C B Desclos, Meghan R Sullivan, Hayley L Rein, Kara A Bernstein, Przemek M Krawczyk, Jean Gautier, Mauro Modesti, Fabio Vanoli, Maria Jasin

## Abstract

Homology-directed repair (HDR), a critical DNA repair pathway in mammalian cells, is complex, leading to multiple outcomes with different impacts on genomic integrity. However, the factors that control these different outcomes are often not well understood. Here we show that SWS1-SWSAP1-SPIDR controls distinct types of HDR. Despite their requirement for stable assembly of RAD51 recombinase at DNA damage sites, these proteins are not essential for intra-chromosomal HDR, providing insight into why patients and mice with mutations are viable. However, SWS1-SWSAP1-SPIDR is critical for inter-homolog HDR, the first mitotic factor identified specifically for this function. Furthermore, SWS1-SWSAP1-SPIDR drives the high level of sister-chromatid exchange, promotes long-range loss of heterozygosity often involved with cancer initiation, and impels the poor growth of BLM helicase-deficient cells. The relevance of these genetic interactions is evident as SWSAP1 loss prolongs *Blm*-mutant embryo survival, suggesting a possible druggable target for the treatment of Bloom syndrome.

## Main

Double-strand breaks (DSBs) are among the most dangerous DNA lesions that can arise in cells and, if not repaired correctly, can lead to genomic instability and tumorigenesis^1^. Homologous recombination, also known as homology-directed repair (HDR), is the predominant pathway to repair DSBs in an error-free manner. The RAD51 recombinase plays a central role, whereby it forms filaments on single-stranded DNA at resected DNA ends through the activity of mediator proteins like BRCA2^2^ and can be stabilized and/or remodeled by RAD51 paralogs, as shown for the yeast and worm proteins^3,4^. RAD51 nucleoprotein filaments subsequently invade a homologous template, typically the sister chromatid, to form a displacement loop (D loop), followed by repair synthesis. DNA helicases can unwind these D loops to promote HDR by the synthesis-dependent strand annealing pathway to result in noncrossovers. Alternatively, D loops that capture the other DNA end can mature into double Holliday junctions, which can be dissolved by BLM, a RECQ helicase deficient in individuals with Bloom syndrome, or resolved by structure-specific nucleases to form crossovers^5,6^. What controls the decision points for these various pathways is not well understood.

SWS1-SWSAP1 is a recently identified complex that is critically important for HDR during meiosis in the mouse, promoting the stable assembly of both RAD51 and DMC1 nucleoprotein filaments at resected DNA ends in spermatocytes^7^. As a result, meiotic chromosomes in *Sws1* and *Swsap1* mutant mice often fail to synapse properly and mice are sterile. SWS1 homologs are readily identified in a number of organisms based on the conservation of the zinc-coordinating domain (Supplementary Fig. 1a)^8,9^. In humans, SWS1 interacts with the SWSAP1 protein, which like RAD51 contains Walker A and B motifs predicted to be important for ATPase activity (Supplementary Fig. 1b)^10^. Additional interactions have also been reported, including with a potential scaffolding protein SPIDR (Supplementary Fig. 1c)^11,12^. SPIDR was initially identified in human cells as a new binding partner of BLM and was reported to act in the same pathway as BLM in Hela cells^11^. SWS1-containing complexes, often termed “Shu” complexes, vary substantially in different organisms with regards to the number and structure of the other protein components, although like SWSAP1 they typically show limited homology to RAD51 and so are considered to be non-canonical RAD51 paralogs (Supplementary Fig. 1d)^13,14^.

**Fig. 1:**
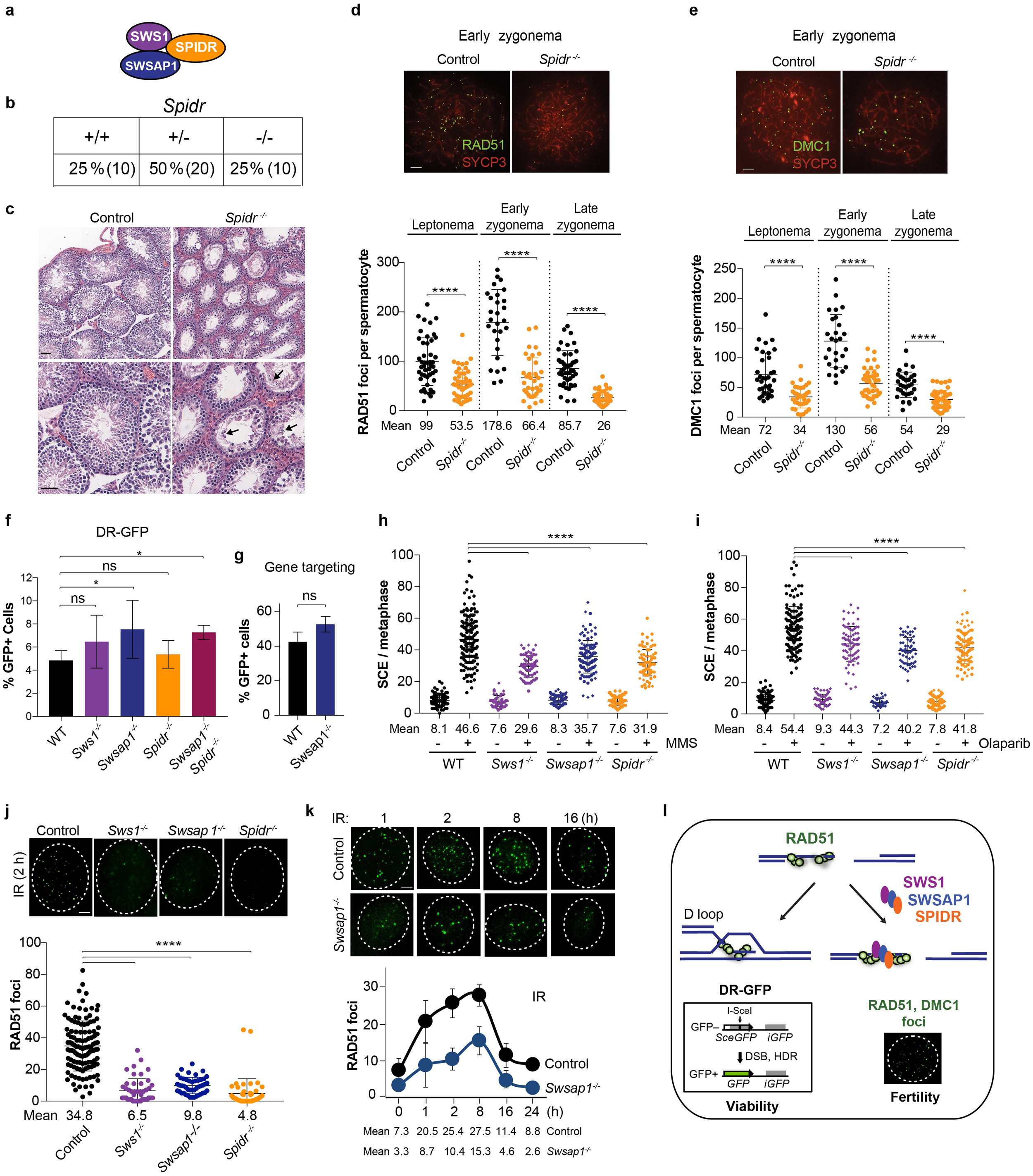
SWS1-SWSAP1-SPIDR promote distinct types of HDR in mitotically-dividing mouse ES cells. **a**. SWS1, SWSAP1 and SPIDR is a three member complex that participates in meiotic HDR and distinct types of mitotic HDR. **b**. *Spidr*^*-/-*^ homozygous mice are viable and born at normal Mendelian ratios. **c**. Testis section from a *Spidr* mutant showing seminiferous tubules with substantially reduced post-meiotic germ cells. Arrows indicate infrequent round and elongating spermatids. Sections were stained with hematoxylin and eosin. Scale bars, 100 μm and 50 μm for top and bottom sections, respectively. **d**,**e**. Representative chromosome spreads from adult mice from early zygonema from control and *Spidr* mutant spermatocytes to analyze RAD51 and DMC1 focus formation. Scale bar, 10 µm. RAD51 (d) and DMC1 (e) focus formation is substantially reduced in *Spidr*^*-/-*^ spermatocytes at early meiotic prophase stages. n=2. (See also **Supplementary Fig. 3c-f**) **f**. HDR levels measured with the DR-GFP reporter are similar in *Sws1*^*-/-*^, *Swsap1*^*-/-*^, *Spidr*^*-/-*^, *Swsap1*^*-/-*^ *Spidr*^*-/-*^, and wild-type ES cells. n≥3. (See also **Supplementary Fig. 5a**) **g**. DSB-induced gene targeting is similar in *Swsap1*^*-/-*^ and wild-type ES cells. Values are 4 days after expression of Cas9-gRNA; in the absence of Cas9-gRNA, GFP+ cells are <0.2%. n=3. **h**,**i**. SCEs are reduced with *Sws1, Swsap1* and *Spidr* mutation in ES cells after treatment with MMS (0.5 mM for 1 h) (h) and olaparib (20 nM for 17 h) (i). Two independent clones are tested for each mutant, as differentiated by diamonds and circles. n≥3. (See also **Supplementary Fig. 5c**,**d**) **j**. *Sws1*^*-/-*^, *Swsap1*^*-/-*^, and *Spidr*^*-/-*^ primary ear fibroblasts have reduced RAD51 focus formation compared to the control cells upon exposure to IR (10 Gy, 2 h). Scale bar, 10 µm. n=3. **k**. Time course of RAD51 focus formation in *Swsap1*^*-/-*^ primary ear fibroblasts has similar kinetics as control cells when treated with IR but with a 2 to 3-fold reduction (10 Gy). Scale bar, 10 µm. n=3. **i**. Efficient formation of observable RAD51 foci is not required for cellular HDR proficiency and hence animal viability, but is required for meiotic recombination and fertility. In our model, the smaller or unstable RAD51 filaments formed in the absence of SWS1-SWSAP1-SPIDR are sufficient for strand invasion and HDR as assayed by the DR-GFP reporter.

Budding and fission yeast Shu complex mutants display a number of HDR-related phenotypes^13^, including relatively mild meiotic defects which can lead to reduced spore viability^15-17^. However, yeast Shu mutants can suppress severe phenotypes associated with deficiency of the DNA helicases Sgs1/Rqh1 (BLM homologs in budding/fission yeasts) and Srs2, for example, the severe DNA damage sensitivity of *sgs1/rqh1* mutants and the *srs2* mutant in fission yeast^8,18,19^. A role for the human SWS1-SWSAP1-SPIDR complex in mitotic HDR has been suggested in studies in human cell lines depleted for this complex, which show reduced RAD51 focus formation and mild sensitivity to DNA damaging agents^8,10,12^. However, the types of HDR promoted by SWS1-SWSAP1-SPIDR and the genetic interaction with BLM have not been delineated. Moreover, the diversity of the Shu complexes in different organisms precludes a direct comparison.

In this study, we show that mouse SWS1, SWSAP1, and SPIDR impact multiple types of HDR. While not required for HDR involving repeats on the same chromosome or gene targeting, these proteins play a crucial role in inter-homolog homologous recombination (IH-HR). This specific defect in IH-HR contrasts with the more general HDR defect observed upon interfering directly with RAD51 function, in which all types of HDR are reduced. Sister chromatid exchanges (SCEs) induced by methyl methanesulfonate (MMS) or inhibition of poly(ADP-ribose) polymerases are partially reduced in *Sws1, Swsap1*, and *Spidr* mutants. More strikingly, SCEs generated by BLM loss are nearly totally abrogated in these mutants, while other HDR events are actually increased with complex disruption. Concomitantly, loss of SWSAP1 in a *Blm* mouse mutant prolongs its embryonic survival. We propose that SWS1-SWSAP1-SPIDR activity plays an important role early in the HDR pathway in determining the fate of the recombination intermediates, where it functions specifically to stabilize intermediates which are processed by BLM.

## Results

### SWSAP1-interacting protein SPIDR is not essential but required during meiosis

Human SWSAP1 has been reported to interact with several proteins in addition to SWS1, including the recently identified scaffolding protein SPIDR^8,10-12,20^. We tested whether the interactions reported for the human SWSAP1 are also present for the mouse protein, confirming that mouse SWSAP1 interacts with SWS1 and SPIDR, consistent with a three member complex SWS1-SWSAP1-SPIDR (Fig. 1a and Supplementary Fig. 1e-g). Mouse SWSAP1 and SPIDR interact with RAD51, and the former also interacts with DMC1, the meiosis-specific RAD51 homolog (Supplementary Fig. 1h-j).

*Sws1*^*-/-*^ and *Swsap1*^*-/-*^ mice are viable^7^, unlike mutants in other HDR genes^21^. Although initially identified as a BLM interacting protein^11^, the interaction with SWSAP1 led us to ask whether loss of SPIDR in mice would phenocopy loss of SWS1-SWSAP1 or whether it is essential like BLM^22^. To test this, we disrupted the gene in fertilized mouse eggs (Supplementary Fig. 2a). As with *Sws1*^*-/-*^ and *Swsap1*^*-/-*^ but unlike *Blm*^*-/-*^ mice, *Spidr*^*-/-*^ mice are born at the expected Mendelian ratio (Fig. 1b) and have a normal adult body weight (Supplementary Fig. 3a,b), demonstrating that SPIDR is not essential in the animal.

**Fig. 2:**
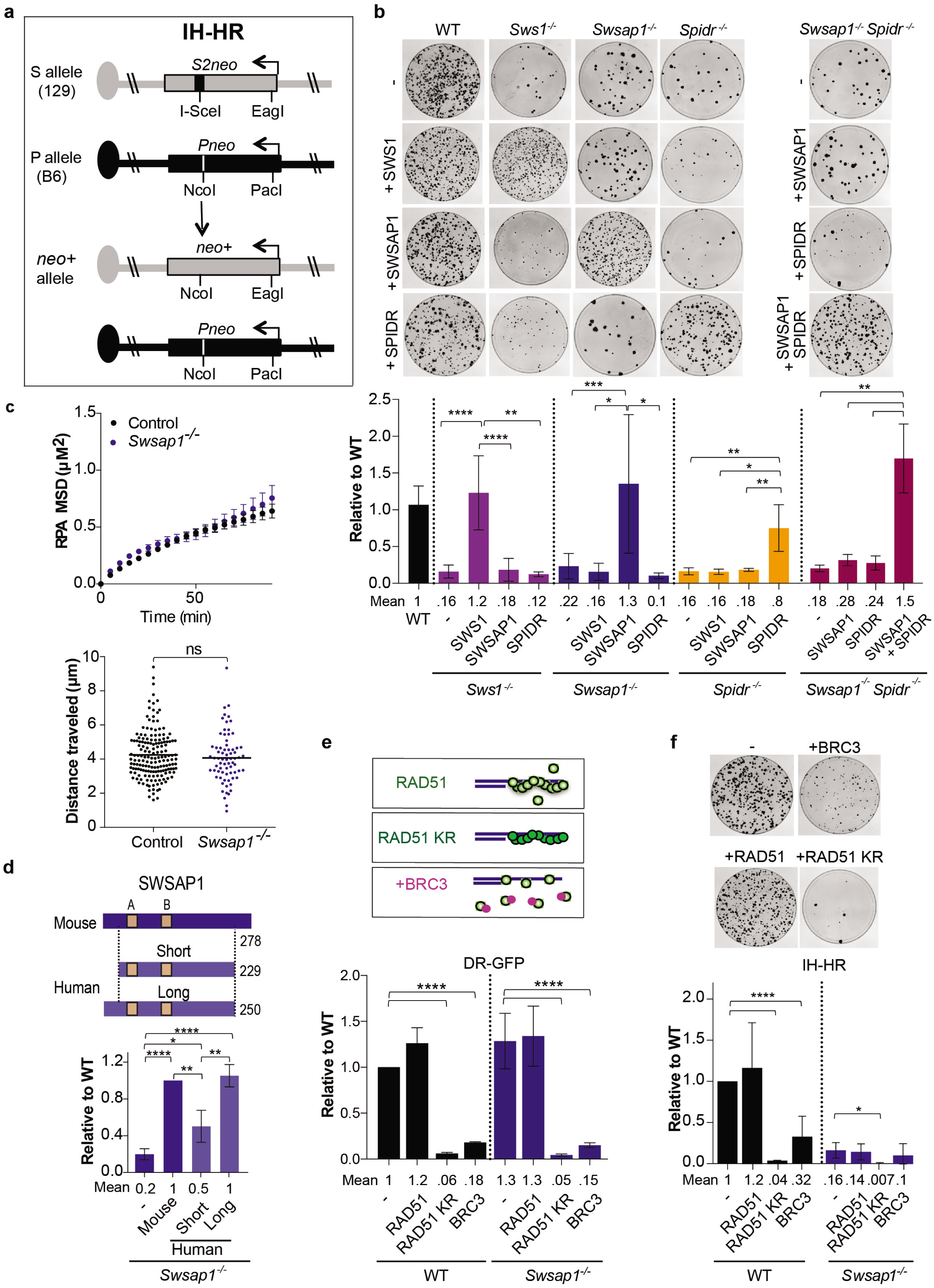
The SWS1-SWSAP1-SPIDR complex promotes IH-HR. **a**. IH-HR at the S/P reporter in 129/B6 hybrid ES cells. The S allele on the 129 chromosome 14 contains the *S2neo* gene mutated by insertion of an I-SceI cleavage site at an NcoI restriction site, whereast the P allele on the B6 chromosome 14 contains the *Pneo* gene mutated by an insertion of a PacI restriction site at an EagI site. After induction of an I-SceI-induced DSB in *S2neo*, IH-HR using the Pneo gene as a template results in a functional neomycin gene (*neo*^+^) that gives rise to G418 resistant colonies. **b**. *Sws1*^*-/-*^, *Swsap1*^*-/-*^, *Spidr*^*-/-*^ and *Swsap1*^*-/-*^ *Spidr*^*-/-*^ cells show decreased numbers of *neo*^+^ colonies, indicative of reduced IH-HR, which is rescued by expression of the cognate proteins. Representative Giemsa-stained plates in S/P reporter cells after I-SceI expression are shown at the top with quantification below. Colony counts here and below are expressed as relative to wild-type for each experiment. n≥3. (See also **Supplementary Fig. 6a**) **c**. Mean square displacement (MSD) analysis of RPA foci following NCS (0.5 μg/mL) treatment in control (*Brca2*^+*/-*^, *Swsap1*^+*/-*^) or *Swsap1*-defective (*Brca2*^+*/-*^, *Swsap1*^*-/-*^) MEFs. n > 450 foci in > 8 cells. Error bars = weighted s.e.m over all MSD curves. Mean cumulative distance traveled by RPA foci following NCS damage in control and *Swsap1*-defective cells is similar. **d**. The human long form of SWSAP1 complements the IH-HR defect of mouse *Swsap1*^*-/-*^ cells to a similar extent as mouse SWSAP1, while the short form has reduced activity. n=4. **e**,**f**. RAD51 K133R (KR) and BRC3 peptide expression reduces HDR in both the DR-GFP (e) and IH-HR (f) reporters in wild-type cells. In *Swsap1*^*-/-*^ cells, RAD51 K133R (KR) and BRC3 peptide expression reduces HDR in DR-GFP to a similar extent as in wild-type cells (e), whereas it further reduces IH-HR (f). RAD51 KR is overexpressed relative to endogenous RAD51 and forms filaments with slow turnover because of the ATP hydrolysis defect. BRC3 peptides can bind RAD51 and sequester it from forming nucleoprotein filaments. n≥3. (See also **Supplementary Fig. 6g**)

**Fig. 3:**
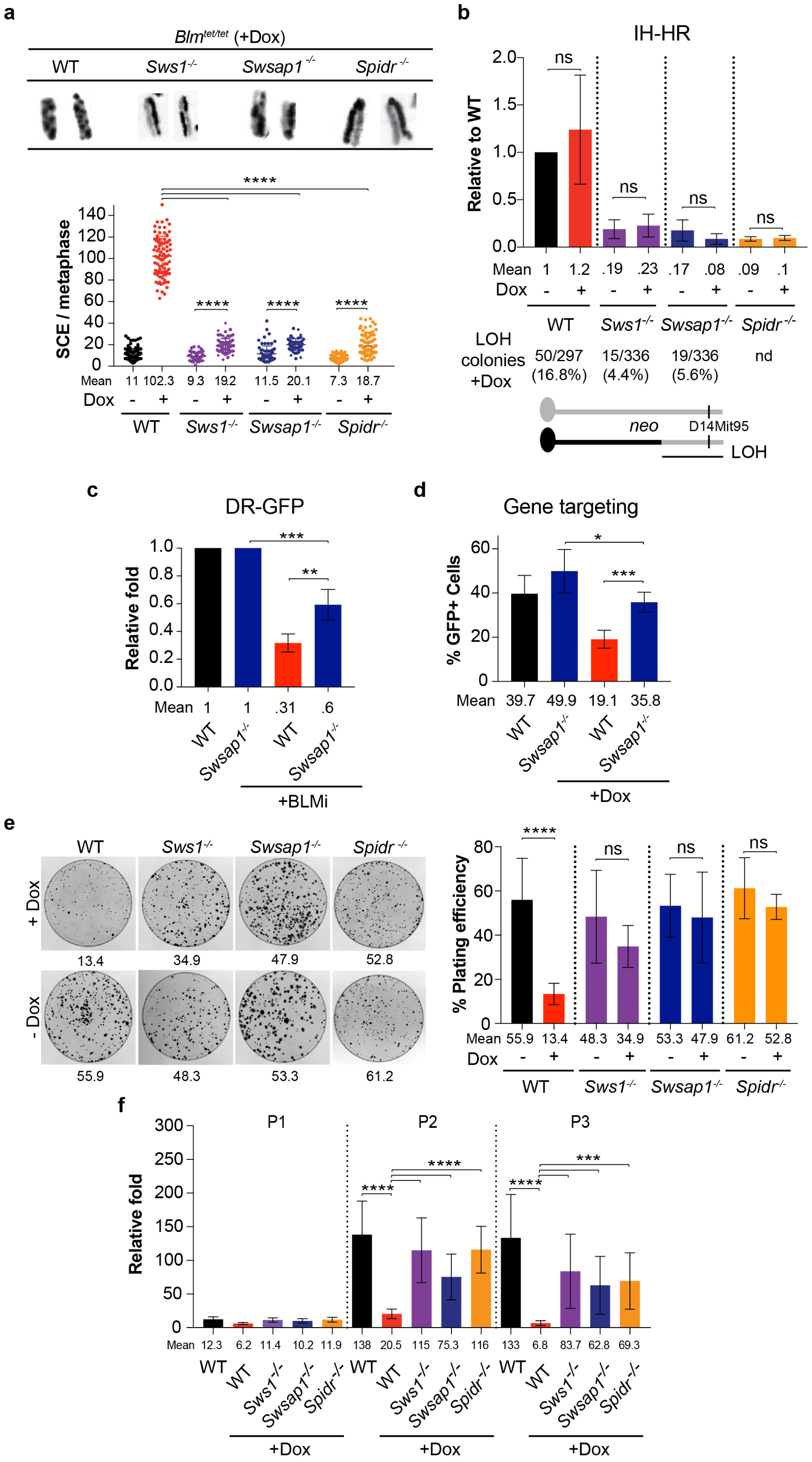
Mitotic phenotypes associated with BLM depletion are dramatically ameliorated by loss of SWS1-SWSAP1-SPIDR. **a**. Loss of SWS1-SWSAP1-SPIDR substantially reduces SCEs in BLM-depleted cells. Representative metaphase chromosomes showing SCEs are shown on the left with quantification upon BLM depletion on the right. *Blm*^*tet/tet*^ cells that otherwise wild type or mutated for *Sws1, Swsap1* or *Spidr* are treated with Dox (1 μM) to deplete BLM. While BLM depletion alone in *Blm*^*tet/tet*^ cells increases SCEs ∼9 fold, BLM depletion in *Sws1*^*-/-*^ *Blm*^*tet/tet*^, *Swsap1*^*-/-*^ *Blm*^*tet/tet*^ or *Spidr*^*-/-*^ *Blm*^*tet/tet*^ cells increases SCEs only by ∼2 fold. n≥3. (See also **Supplementary Fig. 7a**) **b**. IH-HR in SWS1-SWSAP1-SPIDR mutant cells is not further decreased upon BLM depletion, however, LOH of a marker distal to the *neo* locus (D14Mit95) is reduced (p<0.0001), presumably due to reduced crossing over. nd, not determined. (See also **Supplementary Fig. 7b**) **c**. BLM inhibition in wild-type cells reduces HDR in the DR-GFP reporter, however, HDR is partially restored in *Swsap1*^*-/-*^ cells treated with the inhibitor. (BLMi: ML216, 50 uM). n=4. **d**. BLM depletion in *Blm*^*tet/tet*^ cells reduces DSB-induced gene targeting at the *Hsp90* locus, however, gene targeting is partially restored in *Swsap1*^*-/-*^ *Blm*^*tet/tet*^ cells. n=5. **e**. Colony formation is reduced by 3 to 4-fold upon BLM depletion; however, it is restored in BLM-depleted *Sws1*^*-/-*^, *Swsap1*^*-/-*^ and *Spidr*^*-/-*^ cells. The numbers shown below each plate indicates the plating efficiency (%). n=8. (See also **Supplementary Fig. 7c**) **f**. Population doubling experiments shows reduced cell numbers with BLM depletion at passages 2 and 3, which can be significantly restored in the absence of SWS1, SWSAP1, and SPIDR. n=6. (See also **Supplementary Fig. 7d**)

The most pronounced phenotype of *Sws1*^*-/-*^ and *Swsap1*^*-/-*^ mice is the reduced size of their gonads^7^. Similar to these mutants, testes and ovaries of *Spidr*^*-/-*^ mice are about one third the size of control mice (Supplementary Fig. 3a,b). Seminiferous tubules have greatly reduced cellularity due to a paucity of post-meiotic germ cells, indicating defective meiosis (Fig. 1c). Furthermore, as with *Sws1*^*-/-*^ and *Swsap1*^*-/-*^, meiotic RAD51 and DMC1 focus formation is reduced ∼3 fold in *Spidr*^*-/-*^ spermatocytes (Fig. 1d,e and Supplementary Fig. 3c-f). The presence of a few post-meiotic cells, including elongating spermatids (Fig. 1c), indicates a slightly milder meiotic phenotype than *Sws1*^*-/-*^ and *Swsap1*^*-/-*^, although it remains possible that the mutant *Spidr* allele retains some function in the testis. Thus, while SPIDR is not required for mouse embryonic development, it is critical for RAD51 and DMC1 focus formation during spermatogenesis, similar to SWS1 and SWSAP1^7^.

### HDR in mitotic cells does not require efficient RAD51 focus formation

The germ cell-specific phenotypes observed in *Sws1*^*-/-*^, *Swsap1*^*-/-*^, and *Spidr*^*-/-*^ mice led us to question whether SWS1-SWSAP1-SPIDR is required for HDR in mitotic cells, given the embryonic lethality observed for other HDR mutants. To directly measure HDR, we utilized the commonly employed reporter DR-GFP, which measures HDR between direct repeats leading to GFP positive cells following I-SceI endonuclease expression^23^. We first measured HDR in mouse embryonic stem (ES) cells which have a high S phase component and thus may rely heavily on HDR. We disrupted *Sws1, Swsap1*, and *Spidr* in ES cells containing an integrated DR-GFP reporter using CRISPR-Cas9 (Supplementary Figs. 2b and 4a). Testing multiple mutant ES cell lines for each gene, HDR levels are similar in mutant and wild-type cells (Fig. 1f and Supplementary Fig. 5a). *Swsap1*^*-/-*^ *Spidr*^*-/-*^ double mutant cells are also proficient at HDR (Fig. 1f and Supplementary Fig. 5a).

We further quantified HDR in primary ear fibroblasts from *Sws1*^*-/-*^ and *Swsap1*^*-/-*^ mice containing the DR-GFP reporter^24^ and found that HDR levels are similar to controls, as measured directly by the fraction of GFP positive cells or taking into account total I-SceI site-loss (Supplementary Fig. 5b).

HDR was also examined in *Swsap1*^*-/-*^ ES cells using a gene targeting assay. A DSB was introduced at the *Hsp90* locus by CRISPR-Cas9 to induce repair from a promoterless ZsGreen template in a transfected plasmid^25^. As in the direct repeat assay, which measures intrachromosomal HDR, gene targeting levels are also similar in *Swsap1*^*-/-*^ and wild-type cells (Fig. 1g).

Sister chromatid exchanges (SCEs) provide a distinct measure of homologous recombination. In wild-type cells, SCEs are increased substantially by exposure to either methylmethane sulfonate (MMS) or the poly(ADP-ribose) polymerase inhibitor olaparib (∼6 fold for each; Fig. 1h,i). SCEs are also induced in *Sws1*^*-/-*^, *Swsap1*^*-/-*^, and *Spidr*^*-/-*^ cells, but at levels ∼30% lower on average than in wild type (Fig. 1h,i and Supplementary Fig. 5c,d). Thus, these agents induce SCEs in mutant cells but the induction is mildly reduced compared to wild-type cells.

The lack of an observable HDR defect with SWS1-SWSAP1-SPIDR deficiency in some assays (DR-GFP, gene targeting) and minimal effect in another (SCE) is in line with the viability of mutant mice, but is surprising given reports from knockdown experiments in human cell lines that depletion of these proteins reduces RAD51 focus formation following exposure to DNA damaging agents. To determine whether RAD51 foci are reduced in mouse cells, we examined primary ear fibroblasts from *Sws1*^*-/-*^, *Swsap1*^*-/-*^, and *Spidr*^*-/-*^ mice. In each mutant, a substantial reduction in RAD51 foci is observed following exposure to ionizing radiation (IR) (4-9 fold) (Fig. 1j). RAD51 focus formation is also greatly reduced following exposure to MMS (8-10 fold) (Supplementary Fig. 5e). We also monitored the kinetics of RAD51 focus formation in *Swsap1*^*-/-*^ cells and observed reduced RAD51 foci at each time point tested after IR treatment (Fig. 1k). Thus, efficient formation of observable RAD51 foci is not required for cellular HDR proficiency, indicating that smaller or unstable RAD51 filaments that form in the absence of SWS1-SWSAP1-SPIDR are sufficient (Fig. 1l).

### SWS1-SWSAP1-SPIDR promotes efficient inter-homolog recombination (IH-HR)

Thus far, mitotic components specifically required for inter-homolog homologous recombination (IH-HR) have not been identified. Given that SWS1, SWSAP1, and SPIDR are required during meiosis, we considered whether these proteins play a role in IH-HR in mitotic cells, even if their contribution to intrachromosomal HDR appears to be minimal. IH-HR is critical during meiosis, occurring while chromosomes undergo massive structural changes during synapsis, while requiring meiotic-specific proteins like DMC1^26^. By contrast, in mitotic cells, the sister chromatid is the preferred template for HDR. While the homolog can serve as a template for repair in mitotic cells, it is used at low frequency^27^ and in the absence of chromosome synapsis.

To determine whether SWS1-SWSAP1-SPIDR plays a role in IH-HR, we used 129/B6 ES cells carrying the S/P reporter^28,29^. Briefly, two nonfunctional neomycin (*neo*) genes are present at allelic positions on chromosome 14: one is nonfunctional due to the insertion of an I-SceI endonuclease site (S allele) and the other due to the insertion of a PacI restriction site (P allele) (Fig. 2a). Once a DSB is generated by I-SceI endonuclease in the S allele, repair by IH-HR using the P allele as a template results in a *neo*^+^ recombinant. Most IH-HR events are simple gene conversions not involving a crossover, although a fraction are crossovers which are increased by loss of BLM^29^.

We examined S/P reporter cells mutated for each member of the SWS1-SWSAP1-SPIDR complex (Supplementary Figs. 2c and 4b). The recovery of *neo*^+^ recombinants is reduced 5 to 6-fold in each of the *Sws1*^*-/-*^, *Swsap1*^*-/-*^, and *Spidr*^*-/-*^ cell lines relative to wild-type (Fig. 2b and Supplementary Fig. 6a), indicating defective IH-HR. Complementation experiments confirmed that the IH-HR defects are due to loss of the cognate proteins. Similar to single mutant cells, *Swsap1*^*-/-*^ *Spidr*^*-/-*^ cells show a 5-fold reduction in IH-HR, which can be restored only by coexpression of both SWSAP1 and SPIDR (Fig. 2b).

The requirement for SWS1-SWSAP1-SPIDR for IH-HR was unanticipated, given that these proteins are not required for intrachromosomal HDR. After DNA replication, sister chromatids are held in proximity as long as cohesion is maintained; however, homologs are not paired, such that IH-HR may require a more extensive homology search process. We asked, therefore, whether the mobility of DNA ends is affected in cells deficient for SWS1-SWSAP1-SPIDR after DSB formation. Immortalized mouse embryonic fibroblasts (MEFs) were treated with neocarzinostatin (NCS) to induce DSBs, and the movement of GFP-tagged single-stranded binding protein RPA was monitored over time^30^. The mean-square displacement is 4.217 × 10^−5^ μm^2^ s^−1^ in control MEFs and 5.051 × 10^−5^ μm^2^ s^−1^ in *Swsap1*^*-/-*^ MEFs (Fig. 2c), indicating that DNA end mobility is not significantly altered by loss of SWSAP1, and so is unlikely to account for reduced IH-HR.

### SWSAP1 requirements to promote IH-HR

Sequence alignment of mouse and human SWSAP1 reveals two notable differences (Supplementary Fig. 1b): The mouse protein contains a 21-amino acid N-terminal extension, which although not annotated, can also be found encoded within human exon 1 after an alternative translation initiation. In addition, while the human protein has been reported to support ATPase activity involving canonical Walker A and B motifs^10^, the mouse protein lacks critical residues within the Walker A motif (human: GKT vs. mouse: AQT). Given these distinctions, we examined the ability of human SWSAP1 to cross complement the mouse ES cell mutants using the IH-HR assay. While human SWS1 complements *Sws1*^*-/-*^ cells for IH-HR as efficiently as mouse SWS1, the annotated human SWSAP1 – the “short” form – only partially complements *Swsap1*^*-/-*^ cells (Fig. 2d and Supplementary Fig. 6b). However, the “long” form of human SWSAP1 containing the N-terminal extension complements as efficiently as mouse SWSAP1. Thus, the N terminus of SWSAP1 is necessary for full activity, although it does not affect the interaction with either SWS1 or RAD51 (Supplementary Fig. 6c,d).

The canonical RAD51 paralogs have variable requirements for residues involved in ATP binding/hydrolysis, in some cases requiring both the Walker A and B motifs and in other cases only the Walker B motif^21^. To examine the Walker motifs of mouse SWSAP1, we mutated a conserved residue within the Walker B motif (D115A), as well as a non-canonical residue within the Walker A motif (Q37A) and tested them in the IH-HR assay (Supplementary Fig. 6e). Both SWSAP1 Q37A and D115A are able to substantially complement the IH-HR defects of *Swsap1*^*-/-*^ cells, such that only modest defects are observed. Conversely, we also tested the contributions of the human SWSAP1 Walker A and B motifs and found that Walker A (K39A, K39R) and Walker B (D117A) mutants are as active as the wild-type protein (Supplementary Fig. 6f). An exchange of the canonical Walker A motif in the human protein with the non-canonical mouse motif (“Swap”) is also functional. Thus, canonical Walker motifs are apparently not required for SWSAP1 function in HDR.

A RAD51 interaction motif, FxxA, has been identified in several proteins^31,32^, including recently in human SWSAP1 (Supplementary Fig. 1b), such that mutation of this motif from FAAA to EAAE impairs interaction with RAD51^20^. The aromatic ring of the phenylalanine in this motif in a BRCA2 peptide is buried within a hydrophobic cavity of RAD51 in the crystal structure^31^, suggesting that it may be particularly important. We found that mutation to EAAA in either mouse or human SWSAP1 abrogates its ability to rescue the IH-HR defects of *Swsap1*^*-/-*^ cells and to the same extent as mutation to EAAE (Supplementary Fig. 6e,f), highlighting a key role for the phenylalanine within this motif.

### Distinct role of SWS1-SWSAP1-SPIDR compared to core HDR factors

The substantial reduction in IH-HR in *Sws1*^*-/-*^, *Swsap1*^*-/-*^, and *Spidr*^*-/-*^ mutant cells contrasts with the lack of an observable defect in the DR-GFP reporter assay, which involves repair from the sister or same chromatid. We tested whether other factors would show differential effects on HDR in these two contexts. To this end, we expressed a mutant form of RAD51 or a peptide from BRCA2, both of which have previously been shown to reduce HDR in the DR-GFP reporter through different mechanisms, i.e., RAD51 K133R, which forms hyperstable RAD51 nucleoprotein filaments^33,34^, and the BRC3 repeat from BRCA2, which sequesters RAD51 to impede RAD51 filament formation (Fig. 2e)^33,35,36^. RAD51 K133R expression substantially reduces both HDR in the DR-GFP reporter (Fig. 2e and Supplementary Fig. 6g) and IH-HR (Fig. 2f). BRC3 expression also reduces HDR in both assays, although to a lesser extent. Thus, RAD51 K133R or BRC3 expression affects both types of HDR in contrast to the specific reduction of IH-HR in *Sws1*^*-/-*^, *Swsap1*^*-/-*^, and *Spidr*^*-/-*^ cells.

We also tested RAD51 K133R and BRC3 expression in *Swsap1*^*-/-*^ cells and found that HDR in the DR-GFP assay is impacted similarly to that of wild-type cells, consistent with the lack of a role for SWSAP1 in this type of HDR event, even as a “backup” (Fig. 2e and Supplementary Fig. 6g). However, both RAD51 K133R and BRC3 further reduce IH-HR in *Swsap1*^*-/-*^ cells (Fig. 2f). These results further demonstrate a distinct role for SWSAP1, and presumbably the entire complex, in specific types of HDR.

### SWS1-SWSAP1-SPIDR drives sister chromatid exchange and poor growth in the absence of BLM

Cells deficient in the BLM helicase have greatly elevated levels of SCEs^37^ which are dependent on HDR factors^38^. Given that DNA damage-induced SCEs are marginally reduced in the absence of SWS1-SWSAP1-SPIDR, we asked whether SCE induction by genetic perturbation of BLM has similar consequences. In *Blm*^*tet/tet*^ ES cells^39^, depletion of BLM by doxycycline (Dox) addition leads to ∼9-fold induction of SCEs (Fig. 3a and Supplementary Fig. 7a), such that the mean number of SCEs is ∼100 per cell, comparable to what is observed in Bloom syndrome patient cells^40^. Remarkably, BLM depletion in *Sws1*^*-/-*^ *Blm*^*tet/tet*^, *Swsap1*^*-/-*^ *Blm*^*tet/tet*^, or *Spidr*^*-/-*^ *Blm*^*tet/tet*^ ES cells (Supplementary Figs. 2c and 4b) reduces levels of SCEs to nearly normal levels (Fig. 3a and Supplementary Fig. 7a), indicating that SWS1-SWSAP1-SPIDR drives SCE formation in the absence of BLM. The profound reduction in SCEs upon depletion of BLM over the moderate reduction in SCEs by either MMS or olaparib in mutant cells indicates a preference for SWS1-SWSAP1-SPIDR to act in pathways involving distinct types of lesions.

IH-HR in mitotic cells typically leads to only local changes at the site of repair. However, when crossing over occurs during IH-HR, it runs the risk of generating loss of heterozygosity (LOH) from the site of the crossover to the telomere (i.e., long-range LOH), which can contribute to cancer initiation^41,42^. In addition to SCEs, crossing over between homologs is also suppressed by BLM^29^. We found that depletion of BLM does not further decrease IH-HR frequencies in *Sws1*^*-/-*^, *Swsap1*^*-/-*^, or *Spidr*^*-/-*^ cells (Fig. 3b and Supplementary Fig. 7d). However, crossing over upon BLM depletion is reduced by *Sws1* or *Swsap1* mutation, as indicated by fewer colonies with LOH of a distal marker. Thus, deleterious LOH that arises upon IH-HR with BLM depletion is substantially reduced by SWS1-SWSAP1-SPIDR loss by the combined reduction in frequency of IH-HR and fewer LOH outcomes.

To further examine the relationship between BLM and SWS1-SWSAP1-SPIDR in HDR, we used DR-GFP reporter and *Hsp90* gene targeting assays in cells treated with a BLM inhibitor or with Dox to deplete BLM, respectively. With BLM deficiency, we observed a small reduction in HDR in the DR-GFP assay, consistent with previous results^43^, and also in the gene targeting assay (Fig. 3c,d). Notably, concomitant loss of SWSAP1 leads to a partial restoration of HDR in both assays.

BLM has been reported to affect cell proliferation. Primary human and mouse fibroblasts deficient in BLM grow slowly^22,44,45^ and depletion of BLM in *Blm*^*tet/tet*^ ES cells reduces colony formation^29^. Remarkably, loss of SWS1, SWSAP1, or SPIDR significantly rescues colony formation upon BLM depletion, such that the number of colonies and their size are substantially restored (Fig. 3e and Supplementary Fig. 7b). Further, population doubling is also restored. At passages 2 and 3, BLM-depleted ES cells show a substantial reduction in cell number compared to non-depleted cells, whereas depletion of BLM in *Sws1*^*-/-*^ *Blm*^*tet/tet*^, *Swsap1*^*-/-*^ *Blm*^*tet/tet*^, or *Spidr*^*-/-*^ *Blm*^*tet/tet*^ ES cells leads to only a modest reduction in cell number (Fig. 3f and Supplementary Fig. 7c). Thus, the activity of SWS1-SWSAP1-SPIDR not only drives SCEs and LOH in *Blm* mutant cells, but it also interferes with BLM-dependent HDR and impels the slow growth of these cells.

### Loss of SWS1-SWSAP1-SPIDR prolongs embryonic survival of *Blm* mutants

BLM is essential for embryonic development in the mouse^46^. *Blm* mutant embryos are developmentally delayed, show increased apoptosis and severe anemia, and die by embryonic day (E) 13.5^22^. Given that loss of SWS1-SWSAP1-SPIDR restores cell proliferation in the absence of BLM and drastically reduces the high level of SCEs, we asked whether loss of SWSAP1 might also prolong the development of *Blm*^*-/-*^ mice. As expected, *Swsap1*^*-/-*^ animals are born at the expected Mendelian ratio (Supplementary Fig. 7e)^7^ and viable *Blm*^*-/-*^ mice are not obtained. Viable *Swsap1*^*-/-*^ *Blm*^*-/-*^ mice are also not obtained, indicating that *Swsap1* mutation cannot fully rescue the survival of *Blm* mutant mice.

To determine whether SWSAP1 loss could delay the embryonic lethality associated with *Blm* mutation, we performed timed matings and collected embryos at E15.5. At this stage, *Blm*^*-/-*^ mutant embryos are greatly underrepresented from the expected Mendelian ratio (5 fold) (Fig. 4a and Supplementary Fig. 7f), and those few that are obtained are dead, exhibiting developmental delay and severe anemia (Fig. 4b and Supplementary Fig. 8a), as reported. By contrast, *Swsap1*^*-/-*^ *Blm*^*-/-*^ embryos are obtained at nearly the expected ratio, and while a third of these phenocopy the few *Blm*^*-/-*^ embryos that reach E15.5, two thirds are viable at this stage. Although still smaller, these *Swsap1*^*-/-*^ *Blm*^*-/-*^ embryos are more developed, exhibiting hematopoiesis and fully formed digits. We performed a more limited analysis at E12.5 and found that the *Swsap1*^*-/-*^ *Blm*^*-/-*^ embryo obtained at this stage was also more developed than the E12.5 *Blm*^*-/-*^ embryos (Supplementary Fig. 8c). Thus, loss of SWSAP1 can extend embryonic development in the absence of BLM.

**Fig. 4:**
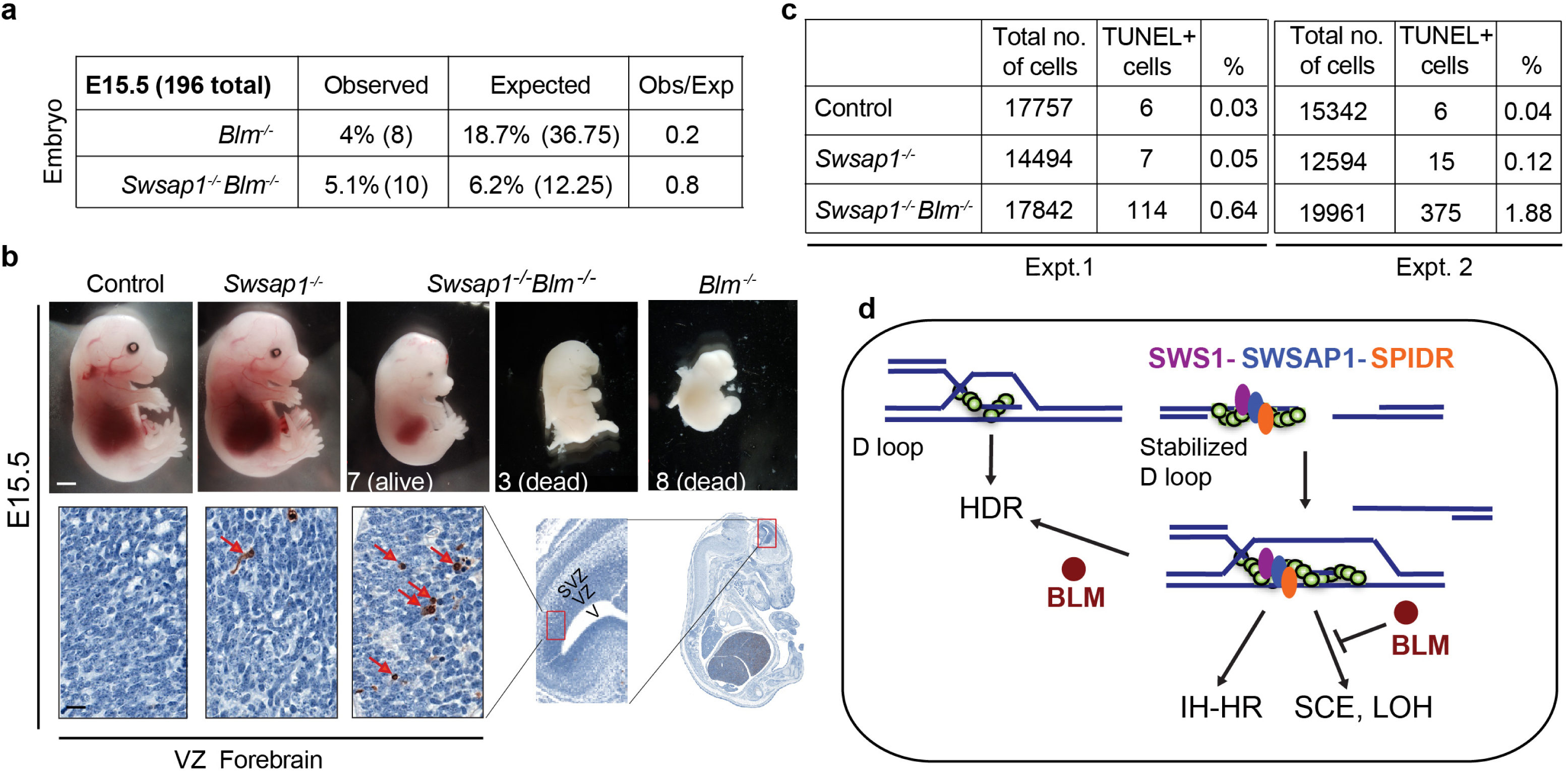
Loss of SWSAP1 prolongs embryonic survival of *Blm* mutants. **a**. *Blm*^*-/-*^ embryos at E15.5 are recovered at 5-fold lower than the expected Mendelian ratio, however, *Swsap1*^*-/-*^ *Blm*^*-/-*^ embryos are recovered only 20% less often than expected. (See also **Supplementary Fig. 7e**) **b**. Analysis of brains from E15.5 embryos. While *Swsap1*^*-/-*^ embryos have normal embryonic development, the few *Blm*^*-/-*^ embryos recovered at this stage are dead. By contrast, most *Swsap1*^*-/-*^ *Blm*^*-/-*^ are alive, but smaller in size, although a few resemble *Blm*^*-/-*^ embryos. Scale bar, 1 mm. Few apoptotic cells are observed in the ventricular zone (VZ) of the forebrain of *Swsap1*^*-/-*^ embryos, while *Swsap1*^*-/-*^ *Blm*^*-/-*^ embryos show numerous TUNEL positive cells (red arrows). The region analyzed in embryos is progressively depicted (red rectangles), with the ventricle (V), ventricular zone (VZ), and sub-ventricular zone (SVZ) indicated. Scale bar, 50 μm. (See also **Supplementary Fig. 8a**,**b**) **c**. Quantification of TUNEL positive cells in the VZ from two experiments. **d**. Model for the role of the SWS1-SWSAP1-SPIDR complex and its genetic interaction with BLM in multiple HDR outcomes. SWS1-SWSAP1-SPIDR functions in the HDR pathway to stabilize RAD51 nucleoprotein filaments which can give rise to stabilized D loops. This stabilization is not required for intrachromosomal HDR through the synthesis-dependent strand annealing pathway, for example, as assayed in the DR-GFP reporter, but it has a major role in promoting IH-HR. These stabilized intermediates are required for efficient formation of dHJs, which can be dissolved by BLM or resolved by strand nicking to generate crossovers and, hence SCEs, when sister chromatids are involved, or LOH when homologs are involved. Thus, in the absence of the SWS1-SWSAP1-SPIDR complex, SCEs and LOH are reduced in the absence of BLM. In addition to dissolution, BLM is known to unwind D loops to promote synthesis-dependent strand annealing; the requirement for this activity may be less in SWS1-SWSAP1-SPIDR mutant cells since RAD51 nucleoprotein filaments are predicted to be inherently less stable, partially restoring HDR, as assayed with the DR-GFP reporter or by gene targeting.

HDR is critical to repair DNA damage arising during neural development, such that in an HDR mutant apoptosis of proliferating cells is observed from early to mid-gestation in the brain^47,48^. *Swsap1*^*-/-*^ embryos have a small increase in apoptosis in the proliferative ventricular zone (VZ) of the forebrain compared to controls at both E12.5 and E15.5 (Fig. 4b,c and Supplementary Fig. 8b,c). By contrast, the viable *Blm*^*-/-*^ embryo obtained at E12.5 had numerous apoptotic cells in its malformed forebrain. *Swsap1*^*-/-*^ *Blm*^*-/-*^ embryos exhibited elevated levels of apoptosis at both stages compared with *Swsap1*^*-/-*^ or control embryos, but not as high as the *Blm*^*-/-*^ embryo. These results may be explained by the partial, but not total, amelioration of SCE and HDR phenotypes in *Blm* mutants by concomitant *Swsap1* mutation.

## Discussion

RAD51 nucleoprotein filaments function at a critical step in the HDR pathway, namely the invasion of homologous sequences which can template repair^49,50^. However, factors that direct these assembled RAD51 nucleoprotein filaments to initiate distinct types of HDR have not been identified. While proteins like BRCA2 that nucleate RAD51 nucleoprotein filaments are apparently critical for all types of HDR, we have uncovered a surprising control of HDR by the SWS1-SWSAP1-SPIDR complex, which contains the non-canonical RAD51 paralog SWSAP1, promoting certain types of events while having little discernible impact on others. A distinct role for the complex clarifies why *Sws1* and *Swsap1* mutant mice^7^, and, as reported here, *Spidr* mutant mice are viable. Loss of these genes in humans is also likely to be compatible with life but lead to infertility. In line with this, two patients homozygous for a *SPIDR* truncation allele have been reported to be developmentally normal except with delayed progression to puberty and ovarian dysgenesis^51^.

Our study has broad impact for understanding the role of factors in HDR, while also providing specific insight into the role of SWS1-SWSAP1-SPIDR. HDR proficiency is typically evaluated in cells by DNA damage-induced RAD51 focus formation and/or by DSB-induced HDR in reporters that measure intra-chromosomal events (DR-GFP). We find that RAD51 focus formation is reduced, but not abolished, in primary mouse fibroblasts deficient for SWS1, SWSAP1 or SPIDR treated with DNA damaging agents, consistent with previous reports in human cell lines^8,10,12^. This parallels what is observed in *Sws1*^*-/-*^, *Swsap1*^*-/-*^, and *Spidr*^*-/-*^ mouse spermatocytes, in which meiotic RAD51 and DMC1 focus formation is also reduced (Fig. 1d,e and Supplementary fig. 3c,d;^7^). By contrast, HDR as assayed between direct repeats, which can reflect HDR between sister chromatids^52^, is not reduced in *Sws1*^*-/-*^, *Swsap1*^*-/-*^, or *Spidr*^*-/-*^ mutant mouse fibroblasts or ES cells.

Unlike for SWS1-SWSAP1-SPIDR, other HDR mutants, e.g., canonical RAD51 paralog mutants, are typically defective in both assays^21,53-56^. While RAD51 focus formation involves a large protein assembly, and so likely has a low threshold for detecting defects, HDR in reporters like DR-GFP typically involves short sequence repeats (<1 kb) and gives rise to short gene conversion tracts (typically <50 bp)^43^, such that only limited RAD51 nucleoprotein filament formation may be required. Given that *Sws1, Swsap1* and *Spidr* mutant mice are viable, while *Brca2* and canonical RAD51 paralog mutants are not^21^, it seems likely that proficient HDR as measured by the DR-GFP reporter reflects physiological intra-chromosomal HDR that is required for survival through embryogenesis, whereas observable RAD51 focus formation is not.

In contrast to intra-chromosomal HDR, SWS1-SWSAP1-SPIDR is important for efficient HDR between homologs, which was completely unanticipated. Although we previously showed that mice defective for SWS1 or SWSAP1 have greatly reduced meiotic IH-HR^7^, it was unexpected that mitotic IH-HR would also need this complex, given that HDR using the DR-GFP reporter or gene targeting is not affected, and moreover that meiotic IH-HR is a highly specialized process. While IH-HR is critical for meiotic chromosome segregation, its role in mitotically dividing cells is more limited and is considered to be potentially deleterious because it can give rise to LOH of tumor suppressor genes^28,29^. These results point to the likelihood of a distinct recombination intermediate during mitotic IH-HR that requires SWS1-SWSAP1-SPIDR but that differs from intermediates during direct-repeat HDR or gene targeting. Because most mitotic IH-HR events are noncrossovers^29^, as are events in the DR-GFP reporter^54^, these intermediates cannot be limited to crossover-bound outcomes.

We also observed a role for SWS1-SWSAP1-SPIDR in promoting SCEs, either DNA damage-induced or arising in the absence of BLM. While the effect is relatively mild for MMS and olaparib-induced SCEs (∼30%), it is enormous in the absence of BLM (5 fold), such that SCEs remain only 2 fold above background levels. Thus, the canonical phenotype of BLM-mutant cells – extremely high SCEs – is largely suppressed by loss of this complex. It is interesting to note that SWS1-SWSAP1 has recently been shown to interact with the cohesion-associated protein PDS5B^12^, which could specifically promote use of the sister chromatid and thus SCE formation. However, a role in determining partner choice during HDR seems unlikely because both IH-HR and SCEs are reduced in the absence of SWS1-SWSAP1-SPIDR. Moreover, mobility of DNA ends is not significantly different in *Swsap1* mutants, as might be expected if cohesion were impaired.

The genetic interactions we observed between SWS1-SWSAP1-SPIDR and BLM are complex: While BLM disruption leads to defects in HDR involving direct repeats, and gene targeting, but greatly increased SCEs and LOH, concomitant loss of SWS1-SWSAP1-SPIDR complex members increases HDR between direct repeats and during gene targeting, while suppressing SCEs and LOH. This complexity of genetic interactions is perhaps not surprising given that BLM has multiple biochemical activities, including in DNA end-resection^57^, D-loop unwinding^58-60^, and double Holliday junction (dHJ) dissolution^61^.

Together, our data suggest a model whereby RAD51 filament stabilization by SWS1-SWSAP1-SPIDR is not required for intrachromosomal HDR, which is considered to be the most common type of HDR and typically involve a synthesis-dependent strand annealing pathway; however, RAD51 filament stabilization by the complex promotes the formation of stable recombination intermediates, including those that are critical for IH-HR (Fig. 4d). Some of these intermediates mature into dHJs which are dissolved by BLM^61^. The absence of SWS1-SWSAP1-SPIDR in *Blm* mutants thus leads to substantially decreased dHJ intermediates and a consequent decrease in SCEs and LOH events.

Consistent with this model, X-shaped recombination intermediates proposed to be dHJs that accumulate in *sgs1* yeast are substantially reduced by additional mutation of Shu complex components^19^.

A model regarding the genetic interaction with SWS1-SWSAP1-SPIDR with BLM also has to take into account a role for BLM in promoting HDR in both the DR-GFP and gene targeting assays and the restoration by loss of the complex. In addition to dHJ dissolution, BLM has also been proposed to act upstream to unwind D loops and promote synthesis-dependent strand annealing (Fig. 4d)^58-60,62^. In our model, the inherently less stable RAD51 filaments and D loops formed in the absence of SWS1-SWSAP1-SPIDR have a reduced requirement for the unwinding activity of BLM, such that loss of SWS1-SWSAP1-SPIDR would help restore HDR in *Blm* mutants. This increase in HDR together with the suppression of SCEs may account for the enhanced cell proliferation in the absence of SWS1-SWSAP1-SPIDR and the prolonged survival of *Blm* mutant embryos when *Swsap1* is also mutated.

Individuals with Bloom syndrome, unlike mice, survive embryogenesis but have developmental issues and a marked susceptibility to various types of cancers^63,64^. Our findings raise the possibility that impairing SWS1-SWSAP1-SPIDR function would suppress symptoms associated with Bloom syndrome. Because *Sws1, Swsap1*, and *Spidr* are not essential genes in mice and at least *SPIDR* appears to be nonessential in humans^51^, interfering with their function may be tolerated in a clinical setting to improve patient outcomes.

## Methods Details

### Immunoprecipitation and western blotting

cDNAs for human and mouse SWS1 and SWSAP1 (with or without an N-terminal FLAG epitope tag), SPIDR, and DMC1 were synthesized by integrated DNA technology (IDT). To clone *Sws1, Swsap1* (wild type and point mutations), *Spidr* and *Dmc1*, pCAGGS was digested with XhoI and KpnI and cDNAs were cloned into it using infusion HD cloning system (Takara Cat # 639645). RAD51, RAD51 K133R, and BRC3 expression vectors in pCAGGS previously described^33^. Synthetic DNA open reading frames coding for human SWSAP1 long and short isoforms and for mouse SWSAP1 with a C-terminal poly-histidine tag were subcloned in frame into the pMBP-parallel vector^65^. The constructs were verified by DNA sequencing. The human RAD51 expression plasmid was previously described^66^.

For immunoprecipitation, 4 µg of the indicated expression vectors were transfected into HEK293 cells using lipofectamine 2000 (Fisher scientific Cat # 11668019). Two days after the transfection, cells were spun at 1000 rpm for 5 min at 4° C. Cell pellets were lysed in NETN buffer (100 mM NaCl, 20 mM Tris-Cl pH 8.0, 0.5 mM EDTA, 0.5% (v/v) Nonidet P-40 (NP-40)) with protease inhibitor cocktail (Roche Cat # 11873580001) by incubating cells on ice for 30 min with gentle flicking every 5 min and then spun at 16000 rpm for 10 min at 4° C to collect the supernatant. Protein concentration was determined using a Bradford protein assay (Bio-Rad). 1 mg extract was used for each immunoprecipitation assay and all the steps were carried out at 4° C. To pre-clear the extract, 25 µl of pre-washed mouse IgG agarose beads (Sigma Cat # A0919) were incubated with 1 mg of extract for 30 min followed by centrifugation at 14000 rpm for 10 min. Supernatant was transferred to a new eppendorf containing 25 µl of anti-FLAG M2 beads (Sigma Cat # A2220) and incubated for 2 h. The anti-FLAG M2 beads with the extracts were spun at 14000 rpm for 3 min, supernatant was discarded, and beads were washed three times with PBST (50 mM Tris [pH 7.5], 150 mM NaCl, 0.05% Tween 20). After the third wash, beads were resuspended in 2x SDS sample buffer (NEB Cat # B7703), boiled for 10 min, and spun at 8000 rpm for 3 min. Supernatant was transferred to a new eppendorf and used for western blots.

To perform western blotting, supernatant was loaded on a precast SDS PAGE gel, transferred onto nitrocellulose membrane, and blocked with 5% milk in PBST (50 mM Tris [pH 7.5], 150 mM NaCl, 0.05% Tween 20) for 1 h. For immunodetection, the following antibodies were used: anti-SWSAP1 (Cat # PA5-25460) from Thermo; anti-RAD51 (Cat # PC-130) from Millipore; anti-DMC1 (Cat # sc-22768) from Santa Cruz Biotechnology; anti-M2-FLAG-HRP (Cat # A8592) from Sigma.

### FRET assay

ARPE-19 retinal pigment epithelial (RPE) cells purchased from ATCC were cultured in DMEM supplemented with 10% FCS and antibiotics. Cells were plated on sterile glass 24 mm coverslips in 6-well culture plates. 24 h later, after reaching 60 % confluence, cells were transiently transfected with 0.5 or 1 µg of DNA per plasmid, when two or one plasmid were used, respectively. Transfections were performed using Genjet transfection reagent. 24 h after transfection, FLIM-FRET (Fluorescence Lifetime Imaging Microscopy-Förster Resonance Energy Transfer) experiments were performed in a 37 °C incubator using a confocal Leica TCS SP8 SMD mounted on a Leica DMI6000 inverted microscope, equipped with a 63x water immersion objective, a white light laser, hybrid detectors, a single molecule detection unit to perform FLIM measurements and a Leica LASX software with FLIM wizard and SymPhoTime64 software. During FLIM measurements, laser intensity was set to the minimal level that produced at most 2×10^6^ counts per second and the repetition rate was set to 20 MHz. Bi-exponential fitting of the fluorescence lifetime data was performed using SymPhoTime64 to obtain amplitude-based average lifetimes.

### Protein purification

Untagged human RAD51 was purified as described^66^. MBP-SWSAP1-His fusion constructs (mouse, human short and human long) were transformed into Rosetta/pLysS bacterial cells (Novagen Cat # 70956) and induced to express the recombinant protein overnight at 15° C after addition of IPTG (2 L scale). Collected cells were lysed in 50 mL of 0,5 M NaCl, 20 mM Tris pH 8, 1 mM EDTA, 1 mM DTT and 10% (V:V) glycerol) supplemented with 1 mM PMSF and protease inhibitor cocktail (Pierce Cat #A32965). The lysate was sonicated and clarified by centrifugation at 20,000 x g for 30 min and the fusion protein recovered by chromatography over 5 ml MPBTrap column (GE Healthcare Cat #28918779) adding 20 mM maltose in the elution buffer. The eluate was next passed through a 5 mL HisTrap Excel column (GE Healthcare Cat #17-3712-06) pre-equilibrated in 0,8 M NaCl, 20 mM Tris pH 8, 1 mM EDTA, 1 mM DTT, 10 mM imidazole and 10% (V:V) glycerol), eluted by raising the imidazole concentration to 0.5 M, aliquoted, snap frozen in liquid nitrogen and stored at −80° C.

### Protein-Protein binding assays

Binding reactions (100 µL) were assembled by mixing RAD51 and various forms of MBP-SWSAP1 each at 0.4 µM final concentration in buffer B (100 mM KCL, 40 mM Tris pH 7.5, 0.05% (V:V) Triton X100 and 100 µg/mL acetylated BSA (Invitrogen Cat #AM2614). Reactions were incubated for 20 min at 37°C in a rotary shaker (600 rpm). Next, the equivalent of 5 µL of magnetic bead slurry (NEB Cat #E8037S, anti-MBP antibody coupled to superparamagnetic beads) passivated with buffer B were added to each reaction and further incubated for 1 h at 37° C. After the reaction, beads were collected with a magnet and washed twice with 100 µL of buffer B and resuspended in 50 µL of Laemmli buffer. The bead pellets (40%) were fractionated by denaturing SDS-PAGE and the gels stained with Coomassie blue or silver for detection of the proteins.

### RAD51 foci formation

For RAD51 focus counts and HDR assays in primary cells, the following mouse alleles were used and genotyped as previously described^7^: *Sws1, Δ1A* and *Swsap1, Δ131*. To prepare ear fibroblasts, ear tips were cut from 2 to 3-month old mice in accordance with IACUC guidelines. Ears were minced with a razor blade and dissociated on a 37°C shaker for 3 h in 3 ml DMEM medium containing 2 mg/ml collagenase A (Roche Cat #11088793001). The dissociated ear fibroblasts were filtered through a 70-μm strainer, and pelleted by centrifugation at 1000 rpm for 5 min. The pellet was resuspended in DMEM medium and plated in a 6-well plate.

Primary ear fibroblasts were expanded and plated in 8-well chamber slides (Thermo Fisher Scientific Cat # C10312) at a density of 20,000 cells per well overnight, followed by 10 Gy IR and 2 h recovery or indicated times of recovery. To label S phase cells, EdU was added right before the IR treatment or 2 h before the cells were fixed on to the slides. To fix, cells were incubated in fresh 3% paraformaldehyde/ 2% sucrose/ PBS at room temperature for 10 min and stored at 4° C. For staining, slides were washed once with PBS, permeabilized in 0.5% Triton X-100/PBS at RT for 10 min, followed by EdU Click-iT (Thermo Fisher Scientific Cat # C10640) to stain S phase cells, and blocked in 1% IgG-free BSA (Jackson ImmunoResearch Cat # 001-000-162) /0.1% Triton X-100/ PBS on a shaker at room temperature for 30 min. Slides were then incubated with RAD51 antibody (1:400; Calbiochem Cat # PC130) for 2 h in blocking buffer, washed with PBS 3 times, followed by incubation with an Alexa Fluor 488-conjugated secondary antibody (Life Technologies, Cat # A21206) for 1 h in blocking buffer, and then washed with PBS 3 times. Slides were fixed in ProLong Gold Antifade Mountant with DAPI (Molecular Probes Cat # P36935).

### DR-GFP and IH-HR assays and LOH analysis

Primary mouse ear fibroblasts were cultured in DMEM-HG medium supplemented with 10% FBS (Thermo Fisher Scientific Cat # SH30070.03), 1X Pen-Strep, 1X MEM/NEAA, and 1X L-Gln. For DR-GFP assays in mouse ear fibroblasts, 0.1 × 10^6^ cells were plated a day before in a 6-well dish, and the next day they were infected with I-SceI lentivirus^67^. 24 h later, virus was removed, cells were washed with PBS, and fresh media was added to plates. The percent GFP^+^ cells were analyzed 48 h later by flow cytometry. Site-loss were determined using the previously described protocol^24^.

All mouse ES cell lines were cultured in DMEM-HG medium supplemented with 12.5% stem cell grade FBS (Gemini), 1X Pen-Strep, 1X MEM/NEAA, 1X L-Gln, 833 U/ml LIF (Gemini Cat # 400-495), and 0.1 mM β-mercaptoethanol. Culture dishes were pre-coated with 0.1% gelatin. For DR-GFP assays, 5 × 10^6^ ES cells were electroporated with 30 µg I-SceI expression plasmid with or without other expression vectors. The trypsinized cells were resuspended in 600 µl Opti-MEM, mixed with plasmids, and pulsed with a Gene Pulser Xcell (Bio-Rad) in a 0.4 cm cuvette at 250 V/950 µF. After pulsing, the cells were mixed with growth medium and plated to a 6-cm dish. The percent GFP^+^ cells were analyzed after 48 h by flow cytometry. For experiments with the BLM inhibitor ML216 (Sigma Cat # SML0661), cells were electroporated with I-SceI expression plasmid and resuspended in growth media containing the inhibitor. The inhibitor was replaced after 24 h and flow cytometry was performed after another 24 h.

For IH-HR assays, 15 x 10^6^ ES cells were electroporated with 30 µg I-SceI expression pCBASce (Addgene 26477) with or without other expression vectors. The trypsinized cells were resuspended in 600 µl PBS, mixed with plasmids, and pulsed with a Gene Pulser Xcell (Bio-Rad) in a 0.4 cm cuvette at 250 V/950 µF. After pulsing, the cells were mixed with 10 ml growth medium and plated to 5 10-cm dishes. 24 h later, one plate was trypsinized to count live cells and the rest of the plates were washed once with PBS and media containing 200 µg/ml G418 (Gemini Cat # 400-111P) was added to select for *neo*^+^ colonies. G418 media was replaced every 3-4 days until the colonies were visible. The plates were fixed in methanol, stained with Giemsa (Life Technologies Cat # 10092-013) and colonies were counted from four plates and the average number of colonies were plotted in graphs. For IH-HR experiments in Fig. 2B, IH-HR is determined by relative fold IH-HR/ plating efficiency. To perform LOH analysis, *neo*+ colonies from each condition were picked into a 96-well plate and cultured till confluency. Cell lysate was prepared using lysis buffer (10 mM Tris pH 8.0, 0.45% NP40, 0.45% Tween 20 and proteinase K 50 µg/ml).The amplification of the distal D14Mit95 marker was done using 1 μl of the lysate as described^29^.

### Gene targeting at the *Hsp90* locus

For gene targeting at the *Hsp90* locus in mouse ES cells, 0.5 × 10^6^ cells were plated in a 6-well dish, and they were transferred to a 10-cm dish the next day. After 48 h, cells were transfected with a plasmid with the targeting vector, a promoterless ZsGreen flanked by homology arms to the *Hsp90* locus, and plasmid DNA for expression of Cas9 and a gRNA directed towards the *Hsp90* locus^25^. Cells were seeded in a 6-well dish after transfection and flow cytometry was performed 4 days later for cells expressing ZsGreen. For experiments where BLM expression was repressed, cells were cultured continuously with 1 µM Dox (Sigma Cat # D9891).

### Sister-chromatid exchange assay

For SCE assays, 0.4 × 10^6^ ES cells were seeded in 6-cm dish and the next day BrdU (10 µM) was added for 24 h for at-least two rounds of DNA replication. For MMS experiments, cells were transiently exposed to MMS (0.5 mM) for 1 h and then were washed with PBS and placed with media containing BrdU. For olaparib treatment, cells were cultured in media containing olaparib (20 nM) and BrdU for ∼17 h. For depletion of BLM, cells were continuously cultured in the presence of 1 µM Dox.

Cells were treated 48 h later in media containing colcemid (0.03 µg/ml) for 40 min. Media was collected in a 15-ml tube, and cells were trypsinized and then collected in the same tube. These cells were centrifuged for 5 min at 1000 rpm and the media was removed. Pre-warmed KCl (75 mM, 4 ml) was added to the pellet and then the tube was inverted gently to resuspend the cells and then incubated for 10 min at 37° C. Five drops (∼100 µl) of cold fixing solution (3:1 v/v methanol-acetic acid) was added to each tube. Tubes were inverted gently and then cells centrifuged for 10 min at 1000 rpm, at room termperature. Media was removed, 5 ml of fixative was added to the pelleted cells; the tube was inverted gently and then incubated on ice for 30 min. Cells were again centrifuged for 10 min at 1000 rpm, at 4° C. These steps were repeated three times and finally cells were resuspended in ∼ 500 µl fixative solution. 10 µl of cells were spotted on a slide from a distance of ∼10 cm. Aged slides were submerged for 45 min in 50 ml of 0.5 x SSC (Saline-sodium citrate) containing Hoechst 33258 (2 µg/ml). Slides were washed for 5 min in 2 x SSC and then were immersed in 50 ml 2 x SSC and exposed for 1 h to UV light. Next, the slides were incubated in 0.5 x SSC at 60° C for 1 h, stained for 15 min in a jar with Giemsa solution in Sorensen phosphate buffer (0.133 M Na_2_HPO_4_, 0.133 M KH_2_PO_4_) at room temperature. The slides were dried at room temperature and then the coverslip was added to image them on a Delta vision ultra high resolution microscope using 100× oil-immersion objective.

### Clonogenic survival and population doubling assays

For clonogenic survival assays, 500 and 1000 cells from IH-HR experiments were plated in a 10-cm plate and were allowed to grow for a week. Plates were then fixed in methanol, stained with Giemsa, and colonies were counted; percent plating efficiency was calculated as the number of colonies divided by the number of plated cells. For population doubling assays, 50,000 cells were plated in 12-well plates at passage 0 and cells were counted every two days until passage 3. For Dox experiments, cells were pre-cultured continuously for 48 h in 1 µM Dox (Sigma Cat # D9891) and during the population doubling assays.

### Live-Cell Imaging and analysis for tracking the movement of GFP-tagged RPA32

Wild-type and mutant mouse embryonic fibroblasts were cultured in high-glucose Dulbecco’s modified Eagle’s medium with L-glutamine, 10% fetal bovine serum, and 1% penicillin-streptomycin. MEFs were plated on 35-mm glass bottom microwell dishes (MatTek Cat #P35GC-1.5-10-C) and transfected with the pEGFP–NLS–RPA32 construct (gift from the Lukas lab) using Lipofectamine 2000 (Invitrogen, 11668). 10 h following transfection, cells were damaged with 0.5 µg/mL neocarzinostatin (Sigma-Aldrich Cat #N9162) for 60 min at 37 °C. Cells were washed twice with PBS and allowed to recover for 14 h prior to imaging. Following recovery, cells were imaged on an A1RMP confocal microscope (Nikon Instruments), on a TiE Eclipse stand equipped with a 60×/1.49 Apo-TIRF oil-immersion objective lens, an automated XY stage, and a stage-mounted piezoelectric focus drive. Live-cell imaging was performed in a heated, humified chamber with 5% atmospheric CO_2_. pEGFP foci were imaged using a 488 nm excitation laser and GFP emission filters. Z-series with 0.4 µm intervals were collected throughout selected nuclei every 5 min for 100 min. Focus was maintained by the Perfect Focus System (Nikon).

DSB foci movement was analyzed as previously described^30^. Briefly, data files exported from Nikon NIS Elements were analyzed in ImageJ^68^, where maximum-intensity z-stack projections were generated. The StackReg plug-in was used to correct for cell movements during the duration of imaging and any nuclei with gross nuclear deformations were discarded^69^. Single-particle tracking was performed using the ImageJ Trackmate plug-in^70^, and trajectories were subsequently analyzed in MATLAB using the @msdanalyzer^71^.

### Yeast-two-hybrid

Yeast-two-hybrid analysis was performed as previously described^12^. pGAD-SWS1 and pGBD-SWSAP1 short or long isoform plasmids were transformed into the *S. cerevisiae* strain PJ69-4A and transformed yeast selected for on SC-LEU-TRP medium. Single transformants were grown overnight in SC-LEU-TRP liquid medium and 5 µl of 0.5 OD_600_ were spotted onto SC-LEU-TRP, SC-LEU-TRP-HIS, SC-LEU-TRP-HIS+3AT, or SC-LEU-TRP-ADE plates. Plates were grown for 2 days at 30° C before being photographed. Images were adjusted identically for brightness and contrast using Adobe Photoshop.

### Generation of *Sws1, Swsap1, and Spidr* mutant cell lines and *Spidr* mice

A dual gRNA approach was used to knockout *Sws1, Swsap1* and *Spidr* in the following ES cell lines: J1-DR1^72^ and 129/B6 *Blm*^*tet/tet*39^. The gRNA sequences were cloned into the dual Cas9/gRNA expression vector pSpCas9(BB)-2A-Puro (PX459, Addgene 48139) according to published protocols^73^. Multiple colonies were picked. Those that gave PCR products indicating a deletion between the two gRNAs were sequenced to identify out-of-frame mutations. In addition to clones which had lost the wild-type alleles, two clones were isolated which maintained the wild-type sequence as additional controls for the DR-GFP assays.

Genotyping for *Sws1* and *Swsap1* was performed as described^7^; genotyping for *Spidr1* was done using the following PCR primers *Spidr*-F: 5′-CCATGTCAAGTTTCCGAGTCATTC-3′, and *Spidr*-R: 5′-AGCATCCTTAGTATGCATAGATTCTAC-3′ under the following conditions: 95° C, 3 min; 35 cycles of 95° C, 30 s; 60° C, 30 s and 68° C, 1 min; and a final extension of 68° C, 5 min.

PCR products were run on a 2.4% agarose gel to resolve the wild-type band from the mutant band. The wild-type product is 470 bp. To generate *Swsap1 Spidr* double mutants, *Spidr* was knocked out using clone *Swsap1#69* in J1-DR1 and *Swsap1#39* in 129/B6 *Blm*^*tet/tet*^ ES cells.

To generate *Spidr* knockout mice, dual gRNA approach was used as described above for ES cells. C57BL6/J zygotes were electroporated (BioRad GenePulser Xcell) by 7 pulses (30V-3ms pulse, 100ms interval) with a mix containing 100ng/ul HiFi Cas9 nuclease (IDT) and 50 ng/ul of each sgRNA (Millipore-Sigma) for *Spidr*. Electroporated zygotes were transferred the same day into pseudopregnant recipients. 68 founders were born and after genotyping them, a heterozygous mouse with Δ77 was chosen for further analysis.

### Timed matings and embryo analysis

*Blm* mice and genotyping have been previously described^22^. *Swsap1*^+*/-*^ *Blm*^+*/-*^ males were mated with *Swsap1*^+*/-*^ *Blm*^+*/-*^ females and plugs were checked to determine pregnancy start date. Pregnant females were sacrificed on 12^th^ or 15^th^ day. Uterine horns were dissected, embryos were collected, photographed, and then fixed overnight in 4% PFA (paraformaldehyde). Embryos that had a heart beat were marked as alive in the analysis. To perform TUNEL analysis, embryos were fixed in 4% PFA and stained with hematoxylin and TUNEL (Roche, 03333566001 and 11093070910, respectively).

### Histology

Testes were collected from animals between 6-8 weeks of age, fixed in 4% PFA, and sectioned and stained with H&E (Hematoxylin and Eosin). Staging of H&E-stained testes sections was performed as described^74^.

### Spermatocyte chromosome spreads and immunofluorescence

Spermatocytes for surface spreading were prepared from testes of ∼2 month old mice using established methods^75^. The following primary antibodies were used in dilution buffer (0.2% BSA, 0.2% fish gelatin, 0.05% Triton X-100, 1xPBS), with incubation overnight at 4° C: goat anti-SYCP3 (Santa Cruz Biotechnology Cat# sc-20845; 1:200), rabbit anti-RAD51 (Calbiochem Cat# PC130; 1:200), rabbit anti-DMC1 (Santa Cruz Biotechnology Cat# sc-22768; 1:200). This was followed by incubation with the following secondary antibodies at 1:500 dilution for 1 h at 37° C: 488 donkey anti-rabbit (Life Technologies Cat#A21206), donkey 594 anti-goat (Invitrogen Cat# A11058). Cover slips were mounted with ProLong Gold antifade reagent with DAPI. Immunolabeled chromosome spread nuclei were imaged on a Deltavision microscope using 100× oil-immersion objective. Only foci colocalizing with the chromosome axis were counted.

### Animal work

The care and use of mice in this study were performed in accordance with a protocol approved by the Institutional Animal Care and Use Committee (IACUC) at Memorial Sloan Kettering Cancer Center (MSK). Mice were housed under Federal regulations and policies governed by the Animal Welfare Act (AWA) and the Health Research Extension Act of 1985 in the Research Animal Resource Center (RARC) at MSKCC, and was overseen by IACUC.

## Statistical analysis

Statistical analyses were performed using a two-tailed Student’s t-test in GraphPad Prism 7. Error bars, mean ± s.d.; ns, not significant; *P ≤ 0.05; **P ≤ 0.01; ***P ≤ 0.001; ****P ≤ 0.0001.

## Data Availability

All relevant data are included in the Supplementary Data File or are available from the lead contact upon request.

## Acknowledgements

We would like to thank members of the Jasin lab for helpful discusssions, especially Tai-Yuan Yu, Agnieszka Lukaszewicz, and Yufuko Akamatsu, as well as Katia Manova and members of the MSK Molecular Cytology core facility for technical help, Agnel Sfeir for the *Hsp90* targeting plasmids, and Peter Romanienko for generating *Spidr* knockout mice. Core facilities at MSK are supported by a Cancer Center Support Grant (NIH P30 CA008748). Research was supported by NIH F32 GM110978 (R.P.), a Paoli-Calmettes Institute PhD fellowship (F.M.), NIH F31 ES027321 (M.R.S.), NIH R01 ES030335 and ACS Research Scholar Grant 129182-RSG-16-043-01-DMC (K.A.B.), the French National League against Cancer (M.M.), and the MSK Functional Genomics Initiative, NIH R35 GM118175 and R01 CA185660 (M.J.).

## Author contributions

R.P., F.V., and M.J. conceived the project and designed experiments. R.P., T.S., B.T., R.W., and F.V. performed experiments, except *in vitro* protein interactions (F.M.), FRET (E.C.B.D., P.M.K.), 129/B6 *Sws1* and *Swsap1* disruption (M.R.S.), Y2H (H.L.R.), and chromosome mobility (J.A.Z.). R.P., F.V., K.A.B., J.G., M.M., and M.J. provided supervision. R.P., F.V., and M.J wrote the manuscript.

