## Supplementary file for "Distinct pathways of homologous recombination controlled by the SWS1-SWSAP1-SPIDR complex"

**Supplementary Figure 1**

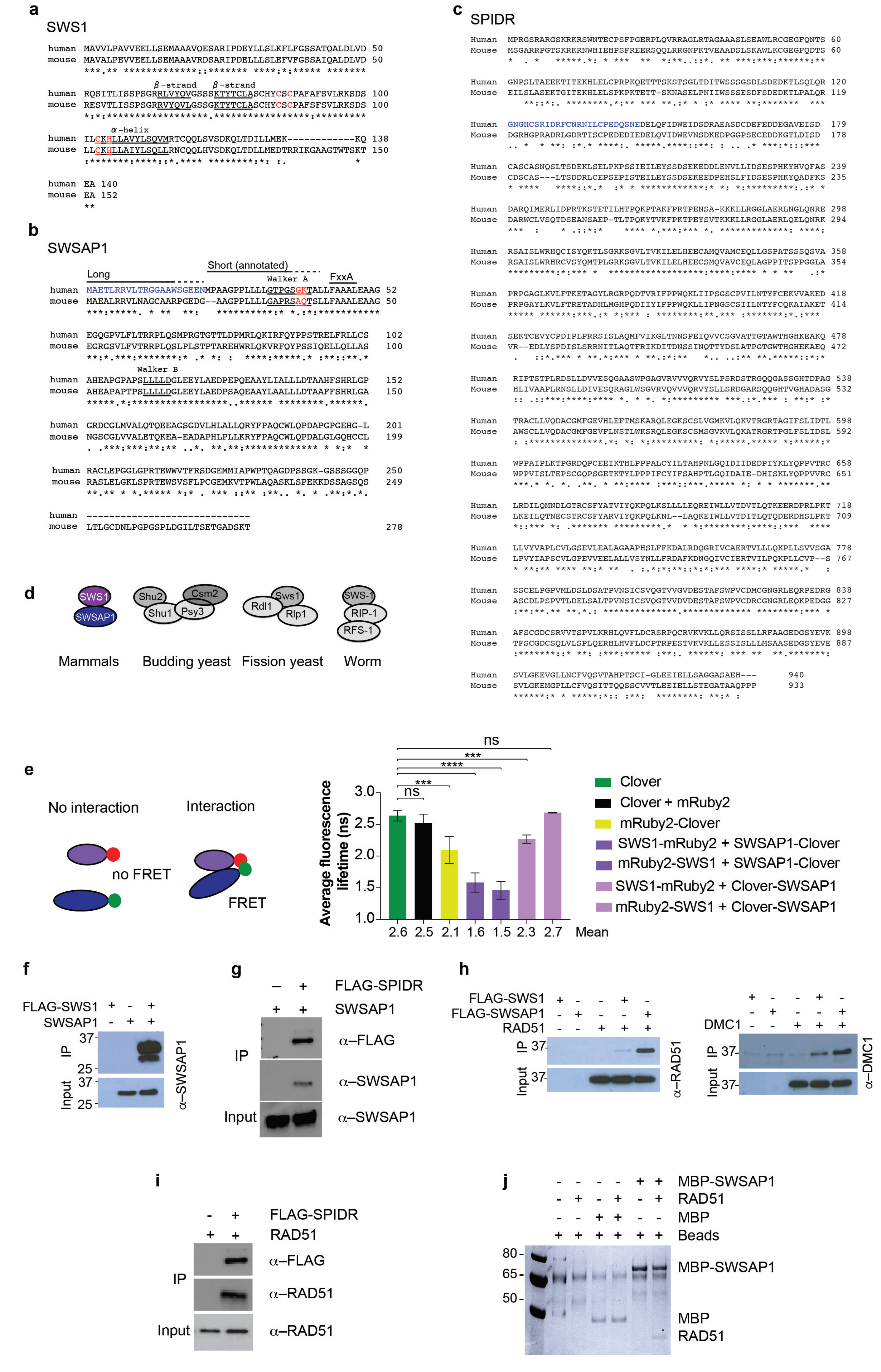

### Supplementary Figure Legends

**Supplementary Fig. 1: Mouse SWS1, SWSAP1, and SPIDR sequences and interactions.**

**a.** Human and mouse SWS1 are 83% identical (96% similar). Each contains a conserved Zn-interacting domain with a CxC($x_{15}$)CxH structure, with the cysteine and histidine residues (red) flanked by two beta-strands and an alpha-helix (underlined) which contain hydrophobic residues characteristic of SWS1 proteins[8,9]. The secondary structure prediction was obtained using YASPIN software. Mouse SWS1 is predicted from the NCBI reference sequence to contain an extra 12 amino acids near the C terminus that were not present in a previous alignment[8]. RT-PCR performed on wild-type testes RNA confirms the presence of the coding sequence for these 12 amino acids.

**b.** Human and mouse SWSAP1 are 69% identical (90% similar). Both share a similar degree of identity with RAD51 (~18%) and are predicted to be RAD51 paralogs[10]. While both the human and mouse proteins contain a conserved Walker B motif (underlined), only the human protein contains a conserved Walker A motif (G$x_4$GKT). The NCBI prediction for human SWSAP1, which was included in[10], has it initiating at the second ATG; however, a start at the first ATG in the transcript gives another 21 amino acids which match well with the mouse and other SWSAP1 proteins (e.g., chimp, macaque) and so we have included it in the alignment. Interestingly, a human *SWSAP1* variant allele disrupts the stop codon to continue translation for another 39 amino acids (WALVGDYLSVLQFPMA GILTSGTYADPKT; not shown), the last 13 of which are very similar to the mouse protein.

**c.** Human and mouse SPIDR are 65% identical (83% similar). The NCBI prediction for mouse SPIDR includes a different splice donor for exon 4 compared with the prediction of the human protein, extending the coding sequence for mouse exon 4 and leading to a gap in the alignment with the human protein (not shown). Using this splice donor for the human protein, however, restores the homologous sequence (shown in blue). Protein alignments were done using ClustalQ program.

**d.** Diversity of Shu complexes in different organisms. A SWS1 homolog is evident in each organism (dark gray), while the number and structure of RAD51-related proteins varies (light gray). See also[13]. The budding yeast complex is the only one for which a structure has been published[14].

**e.** FRET analysis shows that SWS1 and SWSAP1 forms a complex in RPE cells. An interaction with SWS1 is only observed when SWSAP1 is tagged at the C terminus. $n \geq 3$.

**f-i.** Co-immunoprecipitation experiments from HEK293 cells transfected with the indicated expression plasmids. Complex formation is observed between mouse SWS1 and SWSAP1 (f). SPIDR forms a complex with SWSAP1 (g). Interaction for SWS1 and SWSAP1 is observed with RAD51 and DMC1 (h). SPIDR interacts with RAD51 (i).

**j.** Recombinant SWSAP1 binds to RAD51 *in vitro*.

**Supplementary Figure 2**

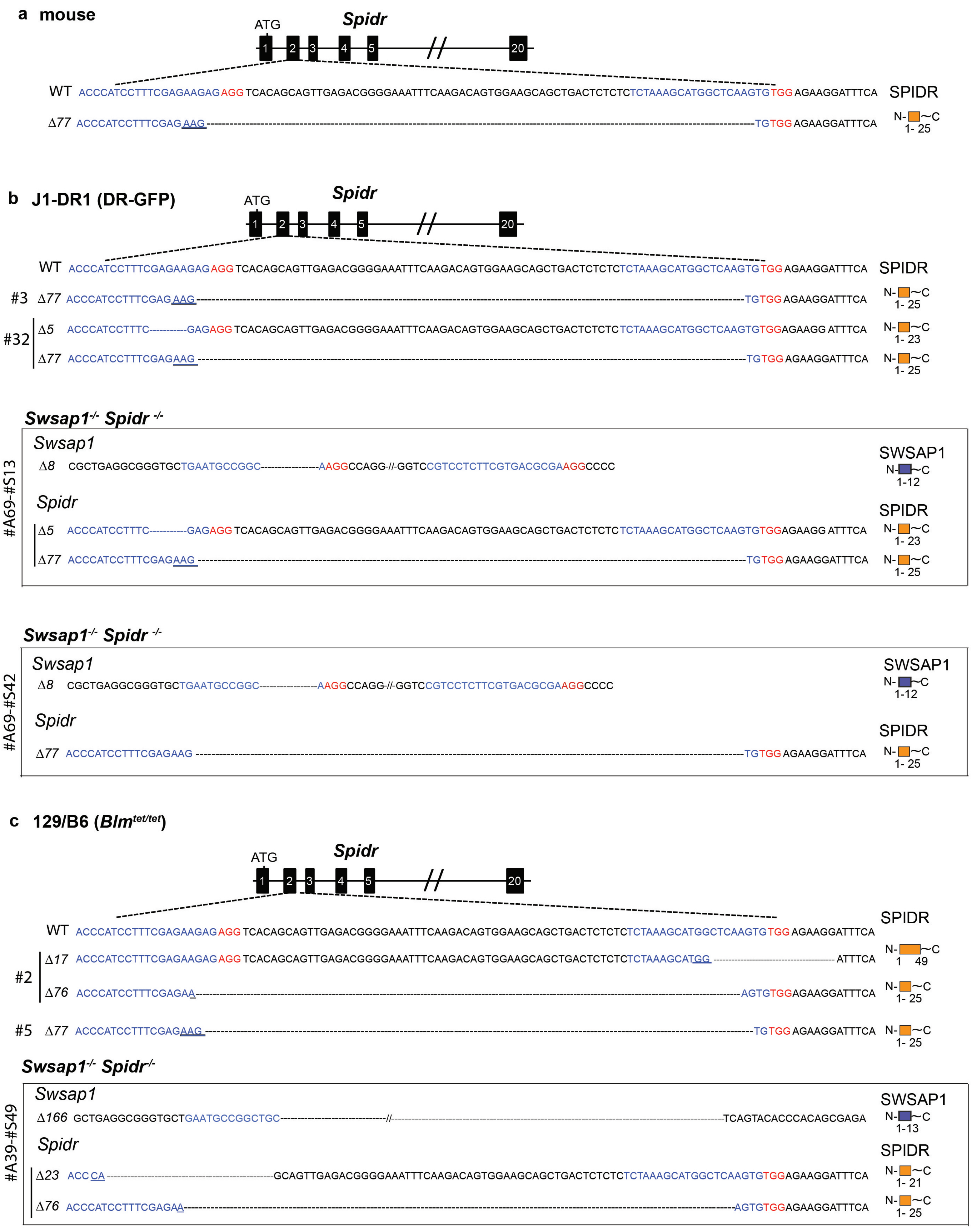

**Supplementary Fig. 2: Mutated alleles analyzed for *Spidr* and *Swsap1 Spidr* in mouse ES cells and mice.**

**a,b,c.** Frameshift mutations in *Spidr* in mice (a), *Spidr* and *Swsap1 Spidr* J1-DR1 (b), and *Spidr* and *Swsap1 Spidr* 129/B6 (c) ES cells. Genomic structure and sequence around the gRNA (blue) and PAM (red) sequences are shown. The number for each mutant clone is shown on the left, together with the detected mutated allele(s). The truncated protein product predicted from each mutated allele is diagrammed on the right.

**Supplementary Figure 3**

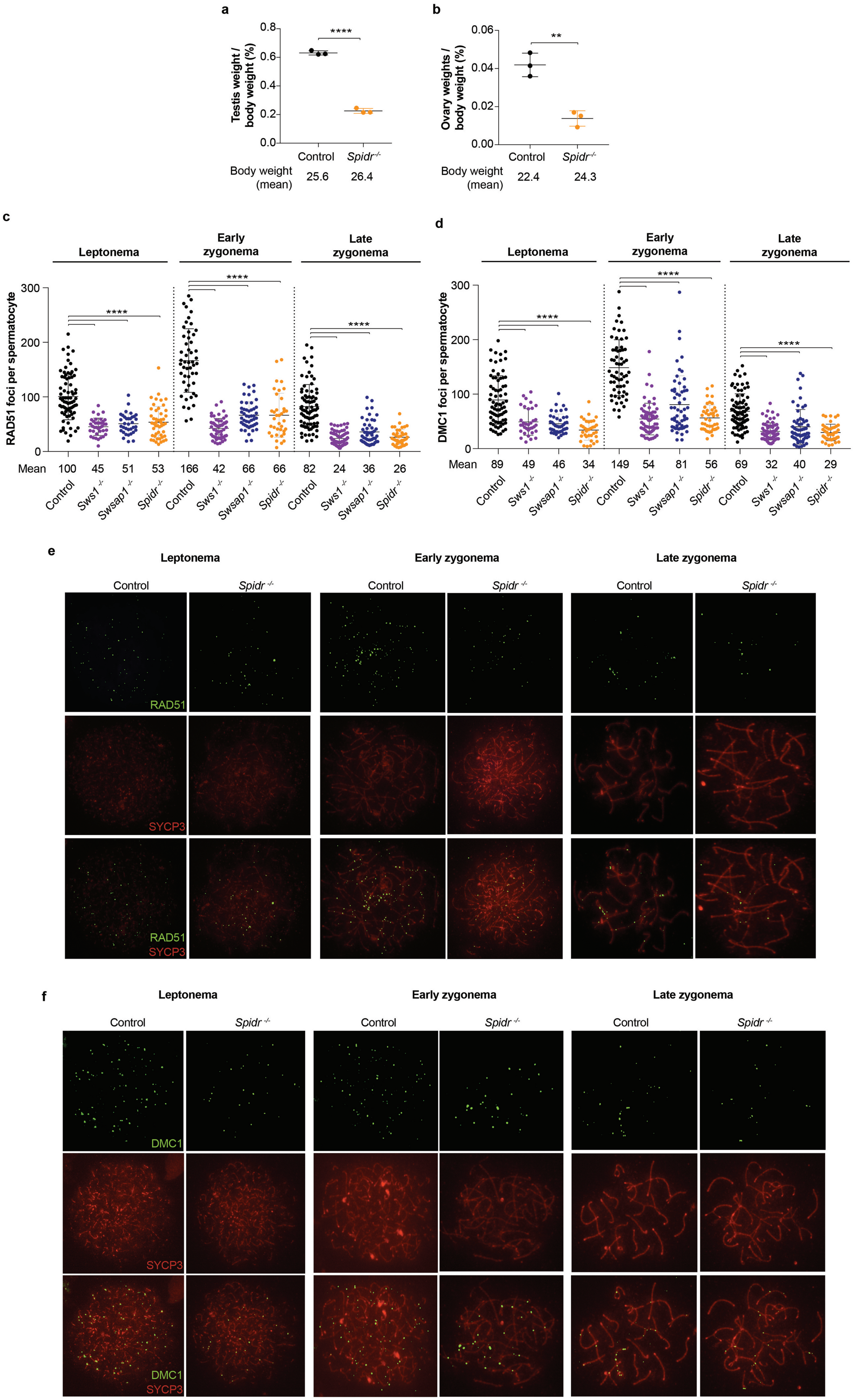

**Supplementary Fig. 3: *Spidr* mutation leads to defect in meiotic HDR.**

**a.** Testes to body weight ratios are significantly reduced in *Spidr*$^{-/-}$ mice. n=3.

**b.** Ovary to body weight ratios are significantly reduced in *Spidr*$^{-/-}$ mice. n=3.

**c,d.** RAD51 (c) and DMC1 (d) foci for data presented in **Fig. 1d,e (**n=2), combined with previously published data from controls, *Sws1,* and *Swsap1*[7]. n= 3.

**e,f.** Representative chromosome spreads from adult mice from various stages of early prophase I from control and *Spidr* mutant spermatocytes to analyze RAD51 (e) and DMC1 (f) focus formation for data presented in **Fig. 1d,e**. Scale bar, 10 μm.

**Supplementary Figure 4**

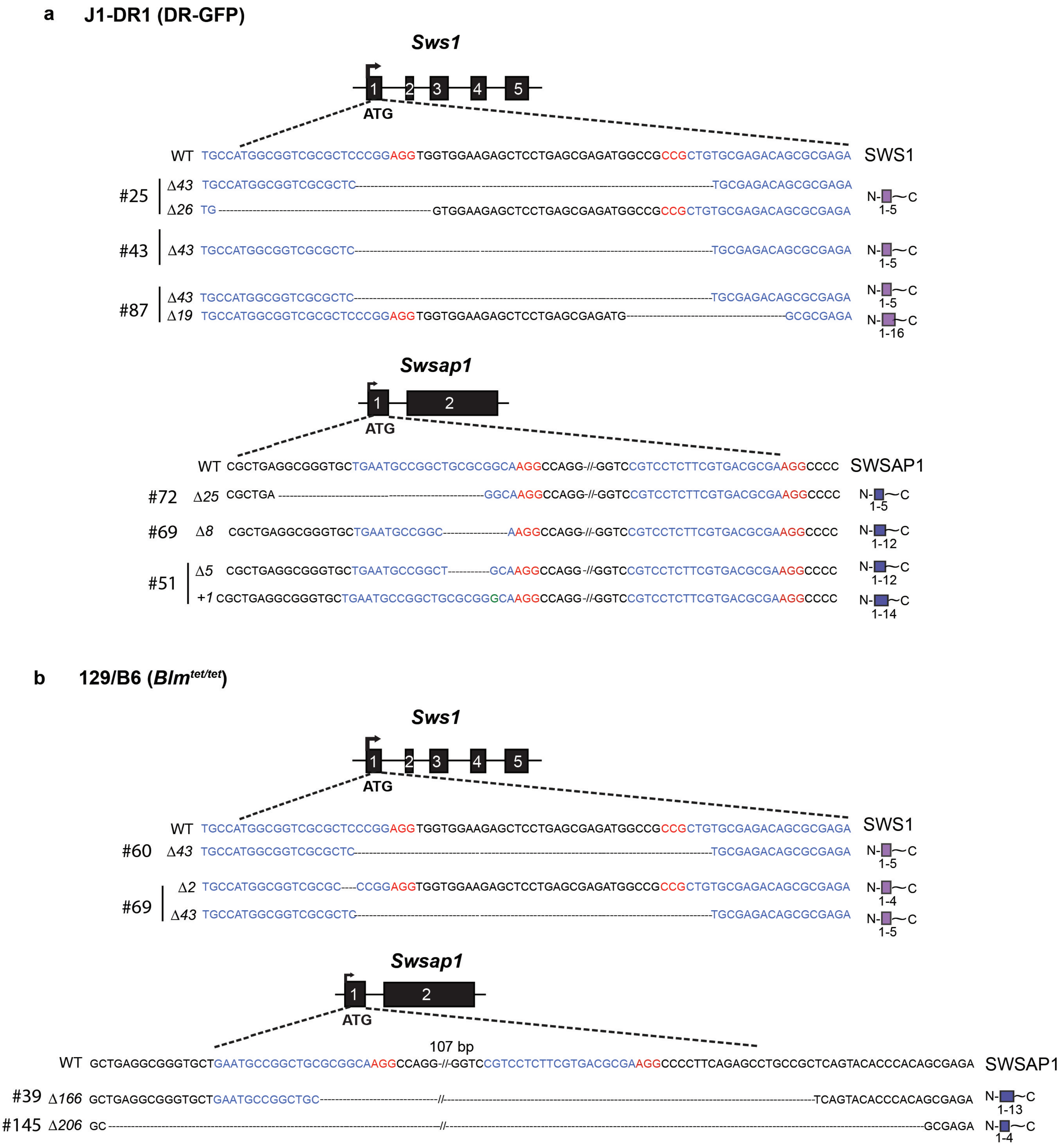

**Supplementary Fig. 4: Mutated *Sws1* and *Swsap1* alleles in mouse ES cells.**

**a,b.** *Sws1* and *Swsap1* frameshift mutations in J1-DR1 (a) and 129/B6 (b) ES cell lines that were analyzed, which contain the DR-GFP and IH-HR reporters, respectively. Genomic structure and sequence around the gRNA (blue) and PAM (red) sequences are shown. The number for each mutant clone is shown on the left, together with the detected mutated allele(s). The truncated protein product predicted from each mutated allele is diagrammed on the right.

### Supplementary Figure 5

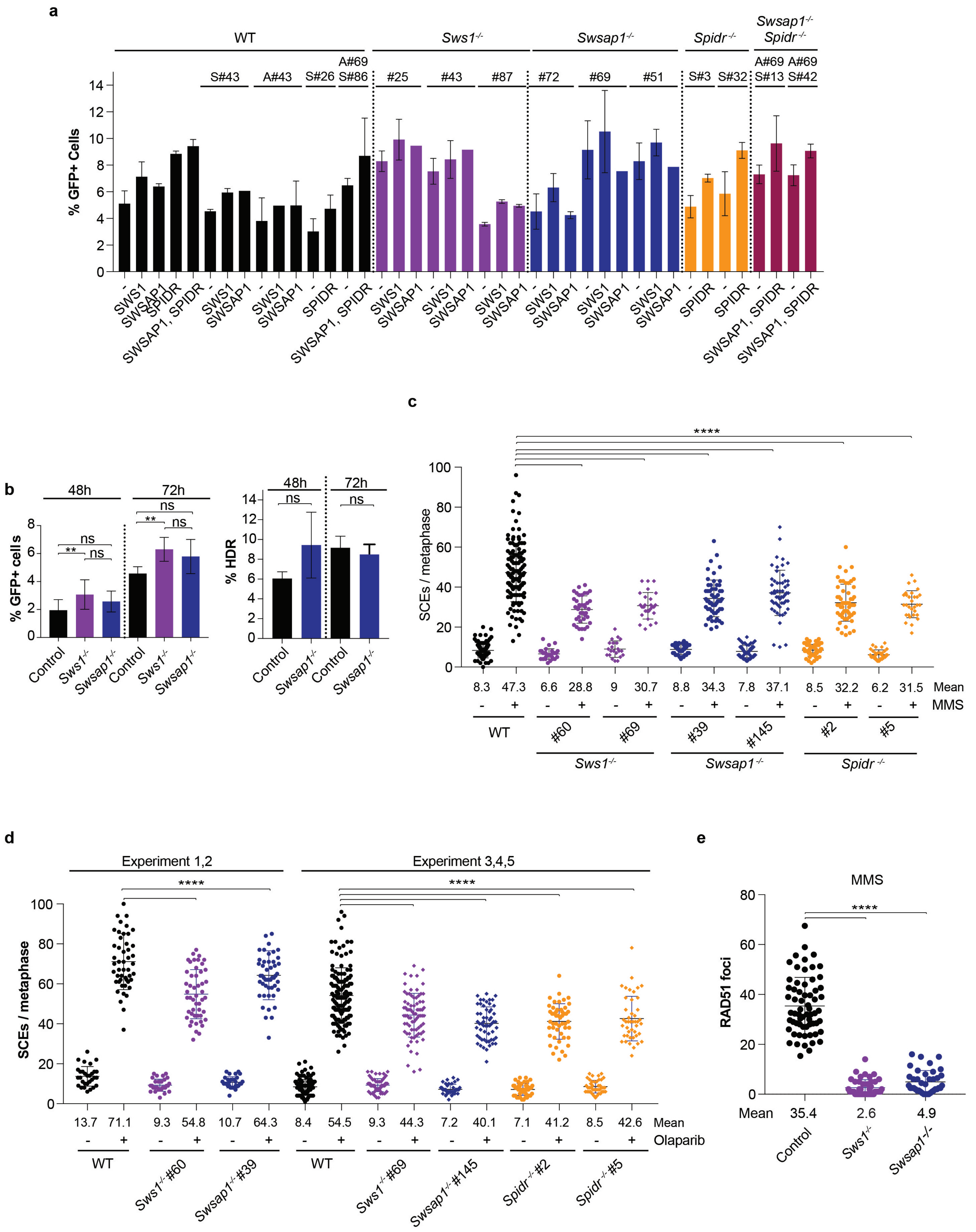

**Supplementary Fig. 5: SWS1-SWSAP1-SPIDR promote distinct types of HDR in mitotically-dividing mouse ES cells.**

**a.** HDR results for individual clones using the DR-GFP reporter in *Sws1*$^{-/-}$, *Swsap1*$^{-/-}$, *Spidr*$^{-/-}$, *Swsap1*$^{-/-}$ *Spidr*$^{-/-}$, and wild-type ES cells for data presented in **Fig. 1f**. expression of SWS1, SWSAP1, SPIDR or SWSAP1 together with SPIDR has no discernible effect. Wild-type includes parental J1 ES cells containing the DR-GFP reporter (J1-DR1) and four subclones isolated after Cas9 expression which maintained wild-type sequences after gRNAs to *Sws1* (S#43), *Swsap1* (A#43), *Spidr* (S#26), *Swsap1* (A#43) *Spidr* (A#69 S#86). n≥3.

**b.** *Sws1*$^{-/-}$and *Swsap1*$^{-/-}$ primary ear fibroblasts have similar level of GFP positive cells as the control at 48 h and 72 h post infection with the I-SceI expression vector. Taking I-SceI-site loss into account to quantify both NHEJ and HDR, the *Swsap1*$^{-/-}$ primary ear fibroblasts have similar HDR levels as control cells. n≥3.

**c,d.** SCEs per metaphase for individual clones for data presented in **Fig. 1h** (c) and **Fig. 1i** (d) in untreated conditions and after exposure to MMS (0.5 mM) and olaparib (20 nM). With olaparib exposure, the mean SCEs was higher in the first two experiments and so results are presented separately from subsequent experiments. n≥3.

**e.** *Sws1*$^{-/-}$ and *Swsap1*$^{-/-}$ primary ear fibroblasts have reduced RAD51 focus formation compared to the control cells upon exposure to MMS (0.5 mM). Scale bar, 10 µm. n=3.

**Supplementary Figure 6**

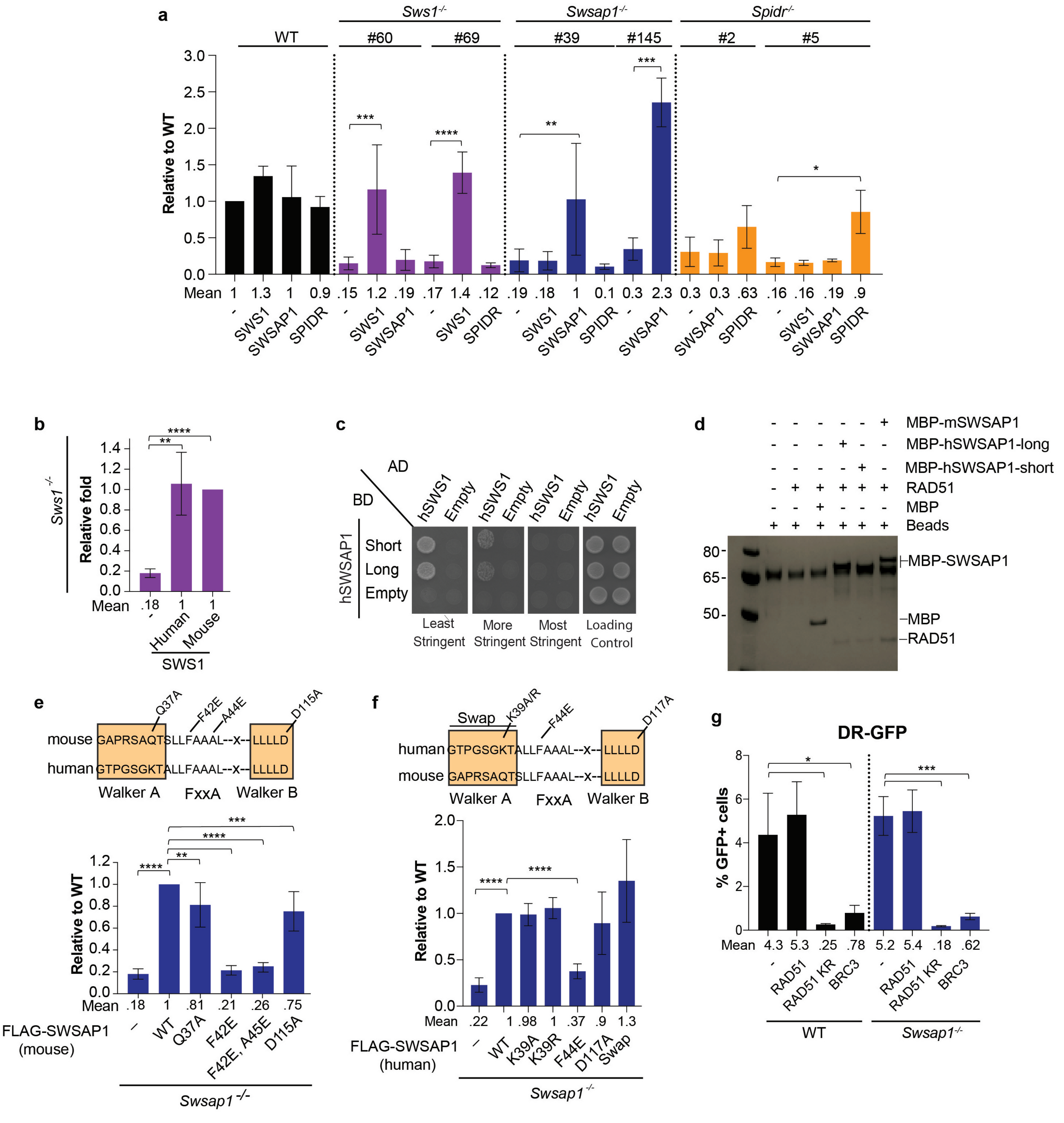

**Supplementary Fig. 6: IH-HR analysis in *Sws1*$^{-/-}$, *Swsap1*$^{-/-}$, and *Spidr*$^{-/-}$ ES cells.**

**a.** IH-HR in individual clones for data presented in **Fig. 2b**. Data are presented as fold relative to wild-type with no expression vector.

**b.** Both human and mouse SWS1 complement the IH-HR defects of mouse *Sws1*$^{-/-}$ cells to a similar level. n=4.

**c.** Yeast-two-hybrid analysis of the short and long forms of human SWSAP1 interaction with human SWS1. Interaction was assessed with SWS1 fused to the GAL4 activation domain (AD) and SWSAP1 short or long form fused to the GAL4-DNA binding domain (BD). Strength of the interaction was assessed by plating on different plates [least stringent (SC-HIS-LEU-TRP), more stringent (SC-HIS-LEU-TRP+3AT), and most stringent (SC-HIS-LEU-TRP-ADE)]. Cells were plated on SC-LEU-TRP as a loading control and the empty vectors were used as negative controls.

**d.** Recombinant long and short isoforms of human SWSAP1 as well as recombinant mouse SWSAP1 bind to RAD51 *in vitro*.

**e,f.** Analysis of mouse (e) and human (f) SWSAP1 mutants for IH-HR activity in mouse *Swsap1*$^{-/-}$ cells. Sequence alignment between mouse and human SWSAP1 shows the conservation in Walker A, Walker B, and RAD51 binding motifs and the residues mutated in this study. Mutations in the Walker motifs (e,f) or "Swap" of the human Walker A motif with the mouse sequence (f) have little or no effect on IH-HR. By contrast, mutation of the phenylalanine in the RAD51 binding motif greatly reduces IH-HR, although additional mutation of the alanine in the FxxA motif has no further effect. e; n≥3. f; n=4.

**g.** Data from **Fig. 2e** shown as percent GFP$^{+}$ cells. n=4.

**Supplementary Figure 7**

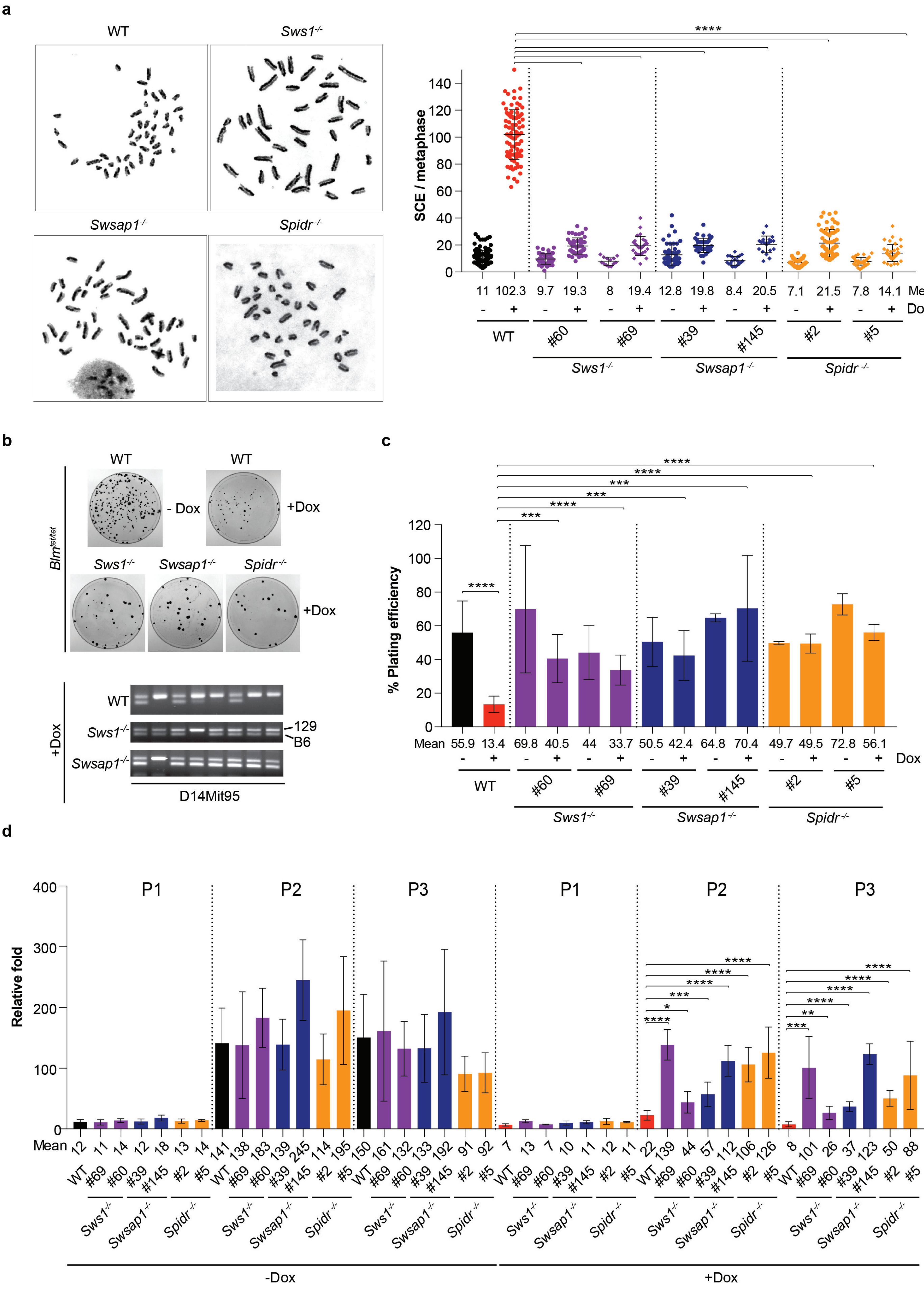


**e**

| Weaning (21d) | Control | | | | Swsap1-/- | | Blm-/- | | Swsap1-/- Blm-/- |
|---|---|---|---|---|---|---|---|---|---|
| **Total (122)** | **wt/wt; +/+** | **wt/wt; +/-** | **Δ/wt; +/+** | **Δ/wt; +/-** | **Δ/Δ; +/+** | **Δ/Δ ; +/-** | **wt/wt ; -/-** | **Δ/wt; -/-** | **Δ/Δ ; -/-** |
| Observed | 7.3% (9) | 17.2% (21) | 14% (17) | 37.7% (46) | 4.1% (5) | 19.7% (24) | 0% (0) | 0% (0) | 0% (0) |
| Expected | 6.25% (7) | 12.50% (15) | 12.50% (15) | 25% (30) | 6.25% (7) | 12.50% (15) | 6.25% (7) | 12.50% (15) | 6.25% (7) |

**f**

| Embryo | Control | | | | Swsap1-/- | | Blm-/- | | Swsap1-/- Blm-/- | |
|---|---|---|---|---|---|---|---|---|---|---|
| **E15.5 (196)** | **wt/wt; +/+** | **wt/wt; +/-** | **Δ/wt; +/+** | **Δ/wt; +/-** | **Δ/Δ ; +/+** | **Δ/Δ ; +/-** | **wt/wt ; -/-** | **Δ/wt; -/-** | **Δ/Δ ; -/-** | **Resorbed** |
| Observed | 9.2% (18) | 18.8% (37) | 9.2% (18) | 31.1% (61) | 6.1% (12) | 16.3% (32) | 1.5% (3) | 2.5% (5) | 5.1% (10) | 18 |
| Expected | 6.25% (12) | 12.50% (24) | 12.50% (24) | 25% (48) | 6.25% (12) | 12.50% (24) | 6.25% (12) | 12.50% (24) | 6.25% (12) | |

**Supplementary Fig. 7: Loss of SWS1-SWSAP1-SPIDR reduces SCEs and LOH, and enhances cell proliferation in the absence of BLM.**

**a.** SCE analysis in *Blm*$^{tet/tet}$ cells after Dox exposure to deplete BLM. Representative metaphase chromosome spreads are shown along with quantification from individual clones for data presented in **Fig. 3a**. n≥3.

**b.** Depletion of BLM in *Blm*$^{tet/tet}$ cells mutated for *Sws1*, *Swsap1* or *Spidr* does not further alter IH-HR frequencies, however, LOH of the distal marker D14Mit95 is reduced. Parental cells are heterozygous for the 129 and B6 alleles of this marker, but *neo*+ colonies that have undergone LOH have a single 129 or B6 allele. Plates and gel images are from experiments in **Fig. 3b**. n=4.

**c.** Plating efficiency for wild-type, *Sws1*$^{-/-}$, *Swsap1*$^{-/-}$, and *Spidr*$^{-/-}$ cells with and without Dox for data presented in **Fig. 3e**. n=8.

**d.** Population doubling for individual clones for data presented in **Fig. 3f**. n=6.

**e.** Breeding analysis showing that *Blm*$^{-/-}$ and *Swsap1*$^{-/-}$ *Blm*$^{-/-}$ mice are not viable upon weaning (21d). Genotypes for *Swsap1* are shown first followed by genotypes for *Blm*.

**f.** Timed matings showing that *Swsap1*$^{-/-}$ *Blm*$^{-/-}$ embryos are obtained at nearly normal Mendelian ratios at E15.5, whereas *Blm*$^{-/-}$ single mutants are highly underrepresented. Partial data from this table are presented in **Fig. 4a**.

### Supplementary Figure 8

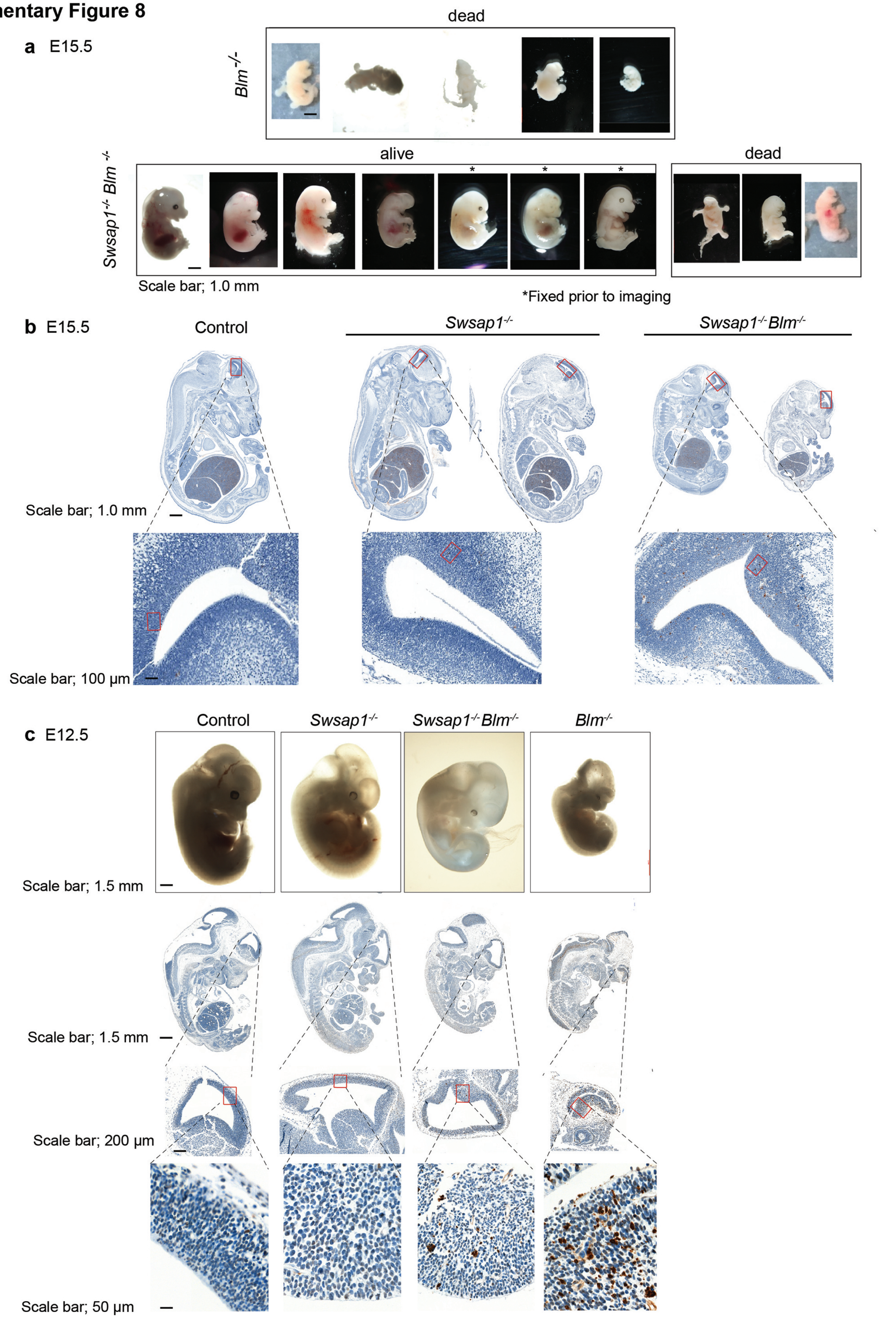

**Supplementary Fig. 8: Loss of SWSAP1 prolongs survival of *Blm* mutant embryos.**

**a.** Images of all E15.5 *Swsap1*$^{-/-}$ *Blm*$^{-/-}$ and *Blm*$^{-/-}$ embryos for data presented in **Fig. 4b**. Scale bar, 1 mm. *Swsap1*$^{-/-}$ *Blm*$^{-/-}$ embryos fixed in PFA for TUNEL analysis before the images were taken are indicated.

**b.** TUNEL-stained sections of E15.5 embryos. Red rectangles in bottom forebrain images show regions in embryo sections that were displayed in lower panels of **Fig. 4b**.

**c.** Images of E12.5 embryos and TUNEL-stained sections. Scale bar, 1.5 mm. The *Swsap1*$^{-/-}$ *Blm*$^{-/-}$ embryo appears better developed compared to the *Blm*$^{-/-}$ embryo and has fewer TUNEL-stained cells, although more than the *Swsap1*$^{-/-}$ and control embryos.
